# Distinct cortical regions support the coding of order across visual and auditory working memory

**DOI:** 10.64898/2026.03.26.714445

**Authors:** Maëliss Vivion, Fabien Mathy, Alessandro Guida, Lydiane Mondot, Stephen Ramanoël

## Abstract

Spatialization in working memory refers to the spatial coding of non-spatial information along a mental horizontal line when encoding verbal material. This phenomenon is thought to support working memory by facilitating order encoding. Although it has been observed for both visually and auditorily presented stimuli, no direct comparison has yet examined whether these modalities rely on similar neural mechanisms. In this study, we investigated whether spatialization in visual and auditory modalities involves shared or distinct patterns of activity within the working-memory network. Forty-nine participants performed both a visual and an auditory working memory SPoARC task of the same verbal material, allowing to study the cortical patterns associated with distinct serial positions at both encoding and recognition across sensory modalities. Whole-brain analyses revealed similar frontoparietal networks across conditions. In addition, a representational similarity analysis (RSA) was conducted to assess the similarity of neural patterns between early and late serial positions in a sequence and across sensory modalities. This multivoxel pattern analysis revealed modality-dependent patterns distinguishing early and late positions in the inferior frontal gyrus. Additional modality-specific effects were observed in the anterior intraparietal sulcus in the visual modality and in the posterior hippocampus in the auditory modality. Drawing on the framework proposed by Bottini & Doeller (2020), we propose that order decoding in the IPS might reflect a low-dimensional spatial coding of order (e.g., along a horizontal axis), whereas order decoding in the hippocampus might reflect higher-dimensional spatial representations or temporal representations.

## Introduction

Serial order working memory, the ability to encode and retrieve a sequence of elements in correct order, is central to various cognitive domains including vocabulary learning (Majerus & Boukebza, 2013; Majerus, Poncelet, Greffe, & Van der Linden, 2006), reading (Laskay-Horváth et al., 2025; Peng et al., 2018), and reasoning (Baddeley, 2003; Kane et al., 2004). However, despite extensive investigation, the exact mechanisms through which order is coded and its associated cortical substrates are still uncertain. Among the hypotheses proposed in the literature, one potential account suggests that order representation is grounded in space (Abrahamse et al., 2014; Guida & Campitelli, 2019; van Dijck & Fias, 2011).

Such spatial grounding can be evidenced through the SPoARC (Spatial-Positional Association of Response Codes, Guida and Lavielle-Guida, 2014) effect, also referred to as the OPE (Ordinal Position Effect, Ginsburg et al., 2014). This phenomenon provides empirical support in favor of a spatial coding of order in working memory by highlighting a link between an item’s serial position within a sequence and its spatial position within a mental space, as measured by response times. For example, in their seminal study, van Dijck and Fias (2011) presented participants with sequences of five items presented in random order (words of fruits and vegetables). After this first memorization phase, participants were asked to perform a categorization task (fruit / vegetable), but only if the item presented was part of the memorized sequence. Participants used lateralized response keys to perform the task (e.g., using their left hand to classify the item as a fruit and their right hand to classify it as a vegetable). The effect commonly observed is that participants tend to respond faster with their left hand to items presented in the beginning of the memorized sequence (e.g., position 1 or 2), and faster with their right hand to items presented in the end of the sequence (e.g., position 4 or 5). This facilitation to respond to first positions with the left hand and to last positions with the right hand suggests that participants internally organized the sequence along a left-to-right mental line (see the SNARC effect for this line of interpretation, Dehaene et al., 1993; Ito and Hatta, 2004; Wood et al., 2008). At the theoretical level, this spatial coding of order has been formally introduced through the mental whiteboard hypothesis (Abrahamse et al., 2014). This hypothesis has been developed within the framework of positional models, which postulate that items are associated with an independent contextual representation of their serial position (i.e., a positional marker), with the specificity that these positional markers are posited to be spatial in nature.

Since the first investigation by van Dijck and Fias (2011), the effect has been largely replicated (e.g., Fenwick et al., 2025; Ginsburg et al., 2014, 2017; Guida, Carnet, et al., 2018; Guida et al., 2016, 2020; Huber et al., 2016; Tian & Fischer-Baum, 2025), with evidence highlighting, for example, the crucial involvement of spatial attention in the effect through specific attentional paradigms (de Belder et al., 2015; van Dijck et al., 2013). However, despite the growing body of behavioral evidence supporting the spatial coding of order in working memory, the mental whiteboard hypothesis remains relatively underexplored, and several outstanding questions remain unresolved. In particular, it is still unsure whether the effect reflects a domain-general mechanism or is restricted to specific types of information.

For instance, when investigating the SPoARC across the verbal, spatial, and visual domains, Ginsburg et al. (2017) could not detect the effect with spatial stimuli, and instead found that the effect was limited to verbal-semantic information. In addition, most SPoARC experiments have been conducted using visual-verbal stimuli, and although the effect has also been observed in an auditory-verbal modality (Guida, Megreya, et al., 2018; Guida et al., 2016), no direct comparison between the two has ever been conducted^1^, leaving unknown whether the spatial coding of order relies on shared mechanisms across sensory modalities. Furthermore, although neuroimaging approaches can provide critical insight into working memory functioning (D’Esposito, 2007; D’Esposito & Postle, 2015), only a limited number of studies have examined the cortical bases of the SPoARC, thus considerably limiting the comprehension of this phenomenon and its generalizability across sensory modalities.

To the best of our knowledge, only two studies have investigated the neural correlates of the SPoARC^2^ (Cristoforetti et al., 2022; Zhou et al., 2021). Both studies reported patterns of results broadly consistent with previous neuroimaging findings using standard working memory tasks. These studies thus revealed a fronto-parietal network, comprising in particular the dorsolateral prefrontal cortex (DLPFC, thought to maintain and manipulate order information, Majerus et al., 2010; Marshuetz et al., 2006; Marshuetz et al., 2000) and the intraparietal sulcus (IPS, associated with attention and sequence processing, Attout et al., 2014; Henson et al., 2000; Majerus et al., 2012; Marshuetz et al., 2000). The involvement of the IPS can be further differentiated between its anterior and posterior subdivisions. The anterior IPS is more directly associated with order coding, whereas the posterior IPS is thought to reflect the engagement of attentional processes in working memory, as it is part of the dorsal attention network (see Majerus, Poncelet, Van der Linden, et al., 2006; Majerus et al., 2007). This attentional involvement has been interpreted as supporting the mental whiteboard hypothesis, which posits the recruitment of spatial attention to support serial order processing (Attout et al., 2014).

This working memory network also comprises the cerebellum (related to visuo-spatial processes, attention, working memory, Strick et al., 2009), the supplementary motor area (SMA, related to temporal processing, or domain general sequence processing, Cona and Semenza, 2017; Protopapa et al., 2019), and the supramarginal gyrus (SMG, related to phonological codes or storing of order information, Majerus, 2019; Papagno et al., 2017). In addition, Zhou et al. (2021) found a specific region in the pre-SMA which correlated with the strength of the SPoARC, suggesting that this region plays a role in spatial-positional associations. Furthermore, although the hippocampus is commonly associated with long-term memory processes, it can also be recruited during working memory tasks (e.g., Attout et al., 2014; Roberts et al., 2018). This region is thought to be involved in order and sequence processing (Hsieh et al., 2014; Long & Kahana, 2019), and has been shown to exhibit distinct activation patterns between the beginning and end of a sequence maintained in working memory (Cristoforetti et al., 2022).

Although consistent with previous working memory findings, the fMRI SPoARC studies by Zhou et al. (2021) and Cristoforetti et al. (2022) have exclusively used visual presentations, thus leaving unknown whether these cortical regions are modality-general or modality-specific. Still, studies of working memory more broadly tend to show that the cortical network described above is modality general. Indeed, activations were consistently observed in the same set of regions when using auditory stimuli (prefrontal and parietal cortices, SMG, SMA; see Arnott et al., 2005; Attout et al., 2019; Kalm and Norris, 2014; Kumar et al., 2016; Majerus et al., 2018; Zhang et al., 2003). These results furthermore tend to be confirmed by an EEG study showing that the fronto-parietal network may operate through shared mechanisms across modalities (Kaminski et al., 2019). In addition to network-level analyses, several studies have also focused on specific brain areas such as the SMG, SMA, or the hippocampus to further examine cross-modal processing. For example, Guidali et al. (2019), showed that repetitive transcranial magnetic stimulation of the SMG impaired order memory similarly across auditory, visuospatial, and motor domains, providing evidence of a cross-modal (or amodal) implication of this region when processing order. A review by Cona and Semenza (2017) has likewise suggested similar functional roles for the SMA across visual and auditory domains. Furthermore, Buzsáki and Tingley (2018) and Buzsáki et al. (2022) proposed that order and sequence processing in the hippocampus is modality independent. These findings overall suggest that both modalities could rely on the same network.

However, studies directly comparing auditory and visual working memory tasks tend to report cortical differences in the involvement of these regions depending on the modality. Using fMRI, Stevens et al. (2000) and Rodriguez-Jimenez et al. (2009) both reported engagement of similar networks, while still observing differences across conditions in terms of amplitude of activation (despite similar performance level across both modalities). Specifically, the auditory modality elicited greater activation in the DLPFC and superior temporal gyrus, whereas the visual modality elicited greater activation in the occipital and cingulate gyri. Crottaz-Herbette et al. (2004) also showed overlap in the fronto-parietal regions activated during visual and auditory working memory tasks, while still showing greater activation in the left dorso-lateral prefrontal cortex for the auditory task, and greater activation in the left IPS for the visual task. These results suggest that, while both modalities rely on the same cortical regions, they might engage these regions in different ways. Accordingly, focusing specifically on activation patterns, Majerus et al. (2018) could not reliably decode attentional load and control across modalities (i.e., training the classifier on one modality and testing it on the other) in the intraparietal cortices, suggesting modality-specific neural patterns in this region. However, in a more recent study using MVPA, Rizza et al. (2024) did not find significant differences in neural patterns between a visual and auditory condition in the fronto-parietal network activated during a working memory task. Thus, it remains uncertain whether visual and auditory order codes share common or distinct patterns of activity within the frontal and parietal regions constituting the working memory network.

### The current study

In this study, we investigated the hypothesis, based on the recent fMRI findings by Rizza et al. (2024), that visual- and auditory-verbal order information is processed using shared, modality-independent order codes within the working memory network. To shed light on this issue, we conducted the first direct comparison between a visual and an auditory spatialization task using functional magnetic resonance imaging. In addition, we aimed to explore the specific representation of different serial positions within a sequence using multi-voxel pattern analyses (MVPA, Haxby et al., 2001). This experiment thus comprised two main aims: First, we aimed to extend prior fMRI evidence on the SPoARC effect (Cristoforetti et al., 2022; Zhou et al., 2021) by testing whether the network supporting order processing during an auditory spatialization task matches the network previously identified with visual stimuli, and whether activity patterns within that network are similar across modalities. Such result would provide evidence that order coding is domain-general rather than modality-specific. We also hypothesized that a specific network related to the processing of item information would be engaged in sensory areas at encoding (Cowan et al., 2011; Majerus et al., 2016, 2018).

Second, we further aimed to assess how order is coded across modalities by investigating whether frontal and parietal regions, as well as the hippocampus, show sensitivity to different serial positions in both the visual and auditory modalities. Specifically, we expected to observe similar patterns between conditions in frontal regions, the intraparietal sulcus and in the hippocampus, while showing discriminability between early and late serial positions within these regions.

## Materials & Methods

### Participants

This study was approved by the Comité de Protection des Personnes (CPP 2022-A02839-34). The experiment was conducted with a sample of 49 students from Université Côte d’Azur (37 women and 12 men, mean age = 20.5 years, *SD* = 2.5, age range = 18 − 30 years). A power analysis conducted using NeuroPower (Durnez et al., 2016) indicated that a minimum of 35 participants was required to achieve sufficient statistical power (> .80). A larger sample size was targeted (N = 50) to align with recent recommendations supporting larger fMRI studies (Geuter et al., 2018), and in anticipation to potential data loss due to head movement in the scanner. All participants were native French speakers, right-handed, and received a monetary compensation in exchange for their participation. Participants had no history of neurological disorder and did not suffer from any visual, auditory, language, motor, memory, or attention disorders. Additionally, they did not report any current use of psychoactive medication. Informed consent was obtained from all participants before the experiment began. One participant was removed from the analysis due to poor hearing during the auditory task. In addition, seven participants were discarded due to excessive movements in the scanner (see fMRI analysis section), resulting in 41 participants included in the analyses.

### Material

In the visual task, the stimuli were displayed on an MRI compatible LCD screen (32-inch, display area 698.4 mm (H) × 392.9 mm (V), resolution 1920×1080, refresh rate 60/120 Hz, NordicNeuroLab, Bergen, Norway). Participants could see the screen using a mirror installed on the head coil. The screen was situated in front of the scanner, approximately one meter away from the participant. In the auditory task, we used NordicNeuroLab MRI-compatible headphones (https://www.nordicneurolab.com) for the transmission of binaural auditory stimuli and noise attenuation (30 dB).

The stimuli used for the spatialization task were a list of ten fruits names (Banana (“banane”), Blueberry (“myrtille”), Cherry (“cerise”), Grape (“raisin”), Kiwi (“kiwi”), Lemon (“citron”), Orange (“orange”), Pomegranate (“grenade”), Raspberry (“framboise”), Watermelon (“pastèque”)). All selected items were bisyllabic words for which the frequency and imageability were controlled based on information collected from the OpenLexicon platform. The stimuli were presented either visually (using colored drawings selected from the IMABASE database; Bonin et al., 2020) or auditorily (with a mean frequency of 182*Hz*, *SD* = 4.3*Hz*).

### Task Description

The experiment comprised eight runs, four runs for the visual condition and four runs for the auditory condition. The conditions were presented in blocks (i.e., starting with the four auditory runs and then performing the four visual runs, counterbalanced between participants), with the order of each run within a condition being pseudorandomly permuted. Each run comprised 16 trials (i.e., 64 trials total per condition), each trial consisting of an encoding phase and a recognition phase.

During the encoding phase, four items were presented sequentially in the center of the screen for 4000 ms with an inter-item interval of 1000 ms (or presented for 1000 ms with an inter-item interval of 4000 ms in the auditory condition). During the recognition phase, two test stimuli to be recognized were presented sequentially, for which participants were asked to produce a yes/no manual response. The recognition phase started by a fixation cross displayed for 2000 ms (or a 440 Hz tone for 1000 ms followed by a blank for 1000 ms in the auditory condition), followed by the presentation of a test stimulus in the center of the screen for 3500 ms (or presented for 1000 ms followed by a blank for 2500 ms in the auditory condition). After 3500 ms, the fixation cross or the tone appeared again, followed by the second test stimulus also presented for 3500 ms (see Figure 1). Participants had to determine for each test stimulus whether it was part of the memorized sequence by pressing either the left index finger button or right index finger button on ergonomic MRI compatible response grips (NordicNeuroLab, Bergen, Norway). For each condition, participants were asked to respond “Yes” using their left hand and “No” using their right hand for two runs, while they were asked to do the opposite for the other two runs. Response time and accuracy was recorded for each response. The inter-sequence interval was variable, following a random Gaussian distribution with a mean of 3000 ms and a standard error of 700 ms.

**Figure 1.**
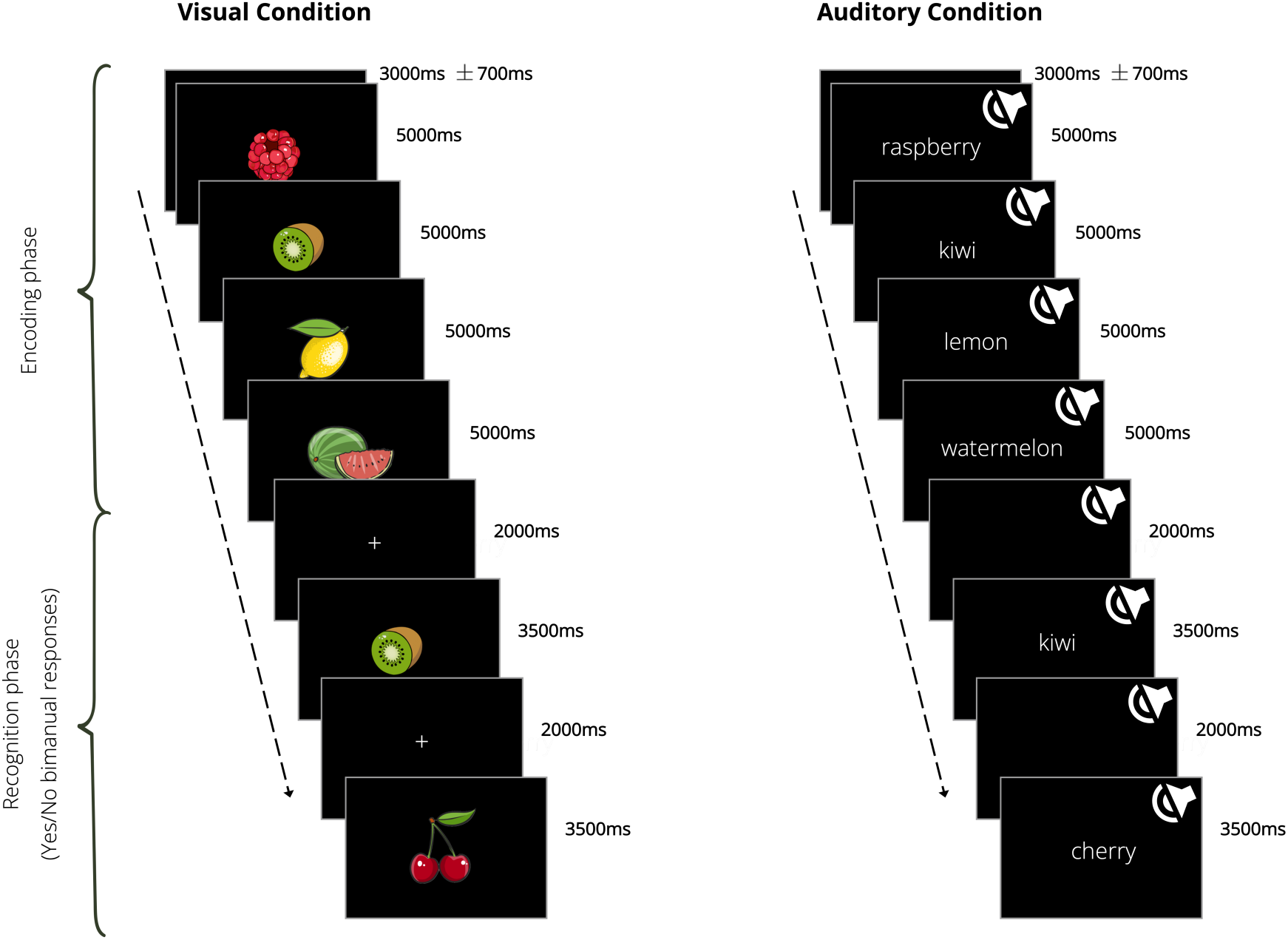
Experimental procedure for a single trial of the visual and auditory task

For both the visual and auditory tasks, the sequences to be memorized were pseudorandomly generated with a controlled number of appearances of each item. For the recognition phase, we ensured that each position in the memorized sequence was equally probed (i.e., 18 probes per position, resulting in 72 probes total per condition). The remaining 56 test stimuli per condition were lures (item not present in the memorized sequence). Before entering the scanner, participants performed four practice trials. The tasks were designed using NordicAktiva 1.3.1 (NordicNeuroLab, Bergen, Norway). The anatomical scan was acquired after the first four runs in order to minimize the temporal gap between the anatomical and functional imaging sessions. The total duration of the experiment was about 1h30.

### MRI acquisition

Imaging data were acquired on a 3T scanner (Discovery, GE Healthcare, Little Chalfont, UK) operated with a 32-channel receiver head coil. Before scanning, foam cushions were used to secure participants’ head and minimize movement. Functional images were acquired using T2*-weighted gradient-echo echo-planar imaging (EPI) sequences covering the whole-brain (TR = 2000ms, TE = 25ms, field of view (FOV) = 236.8×236.8 mm, 64×64 matrix, 38 axial slices with 3.7 thickness acquired along the AC-PC axis, flip angle = 70*^¶^*, voxel size = 3.7×3.7×3.7 mm). Slices were acquired in an interleaved order, with odd-numbered slices acquired first followed by even-numbered slices. For each of the eight runs, 278 volumes were acquired, with the first four scans discarded (dummy scans) to reach steady state of radiofrequency excitation. A high-resolution T1-weighted image was acquired using 3D Axial Brain Volume (BRAVO) sequence for anatomical reference (TR = 8.5ms, TE = 3.2ms, Inversion time = 450ms, FOV = 128×128 mm, 256×256 matrix with 264 slices, flip angle = 12°, voxel size = 0.5×0.5×0.6 mm).

### fMRI analysis

#### Preprocessing

Functional images were preprocessed using SPM12 (Wellcome Centre for Human Neuroimaging, London, UK, http://fil.ion.ucl.ac.uk/spm/) implemented in MatLab R2022b (MathWorks, Natick, USA). First, EPI images were checked for motion artifacts and data with scan-to-scan movements were repaired using an interpolation of the two closest scans. The maximum number of scans repaired was fixed to 3% for each run. As a result, data from 7 participants were discarded due to excessive movement. Across participants, the mean number of scans repaired was 0.18% (*SD* = 0.31%). Next, functional images were corrected for slice timing using the first slice as a temporal reference. Images were then realigned to correct for head motion and distortion, using the default settings in SPM’s Realign module (4th-degree B-spline interpolation), aligning all volumes to the mean image obtained after the first realignment step. This mean functional image was then used to coregister each participant’s structural T1-weighted image using normalized mutual information and 4th-degree B-spline interpolation. Finally, both the structural and functional images were normalized to MNI space (Montreal Neurological Institute). The images produced had a resolution of 3.7mm^3^ (for the functional scan) or 1mm^3^ (for the anatomical scan). Finally, the normalized EPI images were smoothed with an 8 mm full-width half maximum (FWHM) Gaussian kernel for the univariate analyses, and with a 2 mm full-width half maximum (FWHM) Gaussian kernel for the multivariate analyses, in line with recommendations for minimal smoothing in RSA analysis (see Dimsdale-Zucker and Ranganath, 2018).

#### Regions of interest

Regions of interest were defined *a priori* based on previous literature on the SPoARC effect (Cristoforetti et al., 2022; Zhou et al., 2021), and on studies regarding serial order in working memory (Attout et al., 2014, 2019, 2022; Ischebeck et al., 2008; Majerus, Poncelet, Van der Linden, et al., 2006; Majerus et al., 2012; Marshuetz et al., 2006; Marshuetz et al., 2000), because literature regarding the neural bases of the SPoARC is limited.

Eight regions of interest (ROIs) corresponding to both SPoARC and order-related working memory tasks were selected. These ROIs were defined using MarsBaR (Brett et al., 2002) implemented in SPM12. Spheres with a 10 mm radius were centered around the mean coordinates published in the studies previously mentioned (see Table S1 in Supplementary materials for details of the coordinates). First, the intraparietal sulcus (IPS) was considered. This brain region has been extensively associated with order processing in working memory, and activation in this area has been observed during SPoARC tasks. The different subdivisions of the IPS have been linked to distinct processes, with the anterior IPS associated with order processing (Attout et al., 2014), while the posterior IPS is more related to attentional processes (Majerus et al., 2012). Following previous studies, the bilateral IPS was thus divided into two ROIs corresponding to the anterior and posterior IPS (IPSa and IPSp, e.g., Attout et al., 2022; Cristoforetti et al., 2022).

Frontal regions are also known to be involved in order coding in working memory, supporting high-order operations (Amiez & Petrides, 2007). Three frontal ROIs were thus considered: the bilateral inferior frontal gyrus (IFG), the bilateral middle frontal gyrus (MFG), and the bilateral superior frontal gyrus (SFG) (Attout et al., 2014, 2019, 2022; Ischebeck et al., 2008; Majerus, Poncelet, Van der Linden, et al., 2006; Majerus et al., 2012; Marshuetz et al., 2006; Marshuetz et al., 2000). The hippocampus was also included, as it has been suggested to contribute to serial order in memory (Long & Kahana, 2019). Both the bilateral anterior hippocampus (aHC) and posterior hippocampus (pHC) were considered. Finally, the left supplementary motor area (SMA) was included, as this brain region has been shown to be involved in order processing (Cona & Semenza, 2017), and is repeatedly activated in working memory studies (e.g., Attout et al., 2014; Majerus et al., 2016).

#### Univariate analysis

For the first-level analysis, the canonical hemodynamic response function (HRF) in SPM was used to model the blood oxygen level-dependent (BOLD) response in a generalized linear model (GLM). Brain responses were estimated at each voxel using a GLM with both epoch and event-related regressors for each participant. The encoding phase was modeled as an epoch regressor for each modality, ranging from the onset of the first stimulus to the onset of the fixation cross. Two additional event-related regressors were used per modality to model the recognition phase: one regressor for the presentation of a lure and one for the presentation of a probe, with each regressor lasting the duration of the test stimulus. Both conditions were entered into the design matrix, with head motion being accounted for using six regressors (for *x*, *y*, and *z* axis in both translation and rotation). A high-pass filter with a cutoff of 128 seconds was applied to remove low-frequency drifts from the time series. Serial autocorrelations were estimated using a restricted maximum likelihood algorithm with autoregressive modeling (AR1).

Contrasts were formed at the first level, comparing the encoding phase and recognition of a probe of each condition against the implicit baseline. These contrasts allowed us to assess neural activity related to working memory during encoding and retrieval. All contrast images were then entered into a second-level analysis, using paired *t*-tests to assess the significance of the effects. To examine differences between the two modalities, we contrasted conditions during both the encoding phase and the recognition of a probe.

At the whole brain level, activation was estimated at the voxel-level with a *p*-value threshold set to < .05 using a family-wise error correction (FWE) for multiple testing, and a minimal cluster size of *k* = 10 voxels. For the analysis within regions of interest, the mean *β* values within each ROIs were extracted for each participant, and analyses were conducted in RStudio (Posit team, 2023).

#### Multivariate analysis

A representational dissimilarity analysis was conducted to further investigate potential differences in neural response patterns elicited by the different modalities and the different serial positions in the sequence (Kriegeskorte et al., 2008; Popal et al., 2019). This approach allowed us to focus on the activation patterns within our ROIs, rather than on the average voxel activation across the entire region (Dimsdale-Zucker & Ranganath, 2018; Haxby et al., 2001).

For this analysis, the preprocessing steps were identical except that the smoothing was reduced to 2mm (again, in line with recommendations for minimal smoothing in RSA analysis; see Dimsdale-Zucker and Ranganath, 2018). Then, a generalized linear model (GLM) with epoch and event-related regressors was estimated using these minimally smoothed images. This model included 9 regressors per condition: four for the encoding phase (one regressor for each stimulus position) and five for the recognition phase. For the recognition phase, four regressors modeled the recognition of items presented at the first, second, third, and fourth positions, and one regressor modeled the presentation of a lure. As in the first model, six regressors were added to account for head motion (*x*, *y*, and *z* translations and rotations). A high-pass filter with a cutoff of 128 seconds was applied to remove low-frequency drifts from the time series, and serial autocorrelations were estimated using a restricted maximum likelihood algorithm with autoregressive modeling (AR1). Two subsequent analyses were performed, including the regressors corresponding to the encoding and recognition phases, respectively. These analyses were performed using the CoSMoMVPA toolbox (Oosterhof et al., 2016), and custom code in Matlab and Rstudio.

First, the *β* values of activation were extracted from each voxel within our regions of interest (ROIs), providing a mean activation pattern across trials for each regressor and each run. For the analysis concerning the encoding phase, individual representational dissimilarity matrices (RDMs) were computed for each ROI by measuring the distance between activation patterns associated with each pair of regressors (i.e., 1 - *r*, with *r* being the Pearson correlation coefficient between pairs of regressors). The regressors in this analysis corresponded to the encoding of a probe at each of the four probe positions, across the two sensory modalities (visual and auditory), resulting in a total of eight regressors. The resulting matrix was of size 8 × 8 and symmetrical around a diagonal of 0 (as the dissimilarity between each regressor and itself is zero by definition). The same process was applied for the analysis concerning the recognition phase, with the regressors of interest corresponding to the recognition of a probe at each of the four probe positions for each modality, instead of its encoding.

These neural matrices were then compared to theoretical matrices constructed beforehand, which modeled the expected dissimilarity between our regressors from different theoretical perspectives. For each of our two factors (serial position and sensory modality), two possible perspectives were considered, resulting in four theoretical matrices, (see Figure 2): (A) similarity across conditions and distinctiveness between early and late positions (*Across-modality • Early-late position* model); (B) similarity across conditions and distinctiveness between middle and boundary positions (*Across-modality • Boundary-position* model); (C) dissimilarity across conditions and distinctiveness between early and late positions (*Within-modality • Early-late position* model); and (D) dissimilarity across conditions and distinctiveness between middle and boundary positions (*Within-modality • Boundary-position* model).

**Figure 2.**
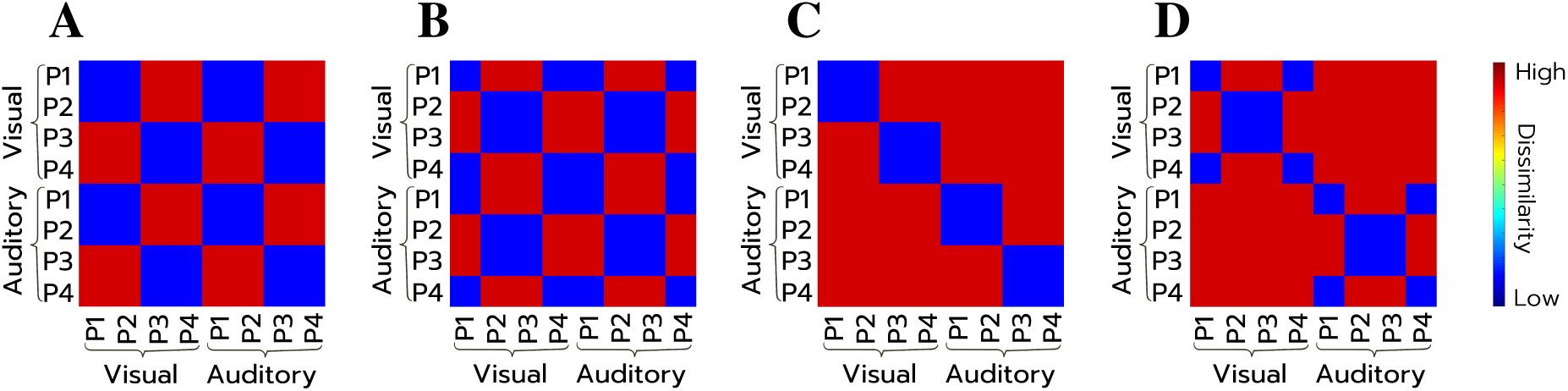
Theoretical representational dissimilarity matrices (RDMs) used in the multi-voxel pattern analysis. **A**, Across-modality • Early-late position model, in which the patterns between modalities are hypothesized to be similar, and early positions (i.e., positions 1 and 2) are hypothesized to be similar to one another but dissimilar from the remaining positions (i.e., positions 3 and 4); **B**, Across-modality • Boundary-position model, in which the patterns between modalities are hypothesized to be similar, and boundary positions (i.e., positions 1 and 4) are hypothesized to be similar to one another but dissimilar from the remaining positions (i.e., positions 2 and 3); **C**, Within-modality • Early-late position model, in which the patterns between modalities are hypothesized to be dissimilar, and early positions (i.e., positions 1 and 2) are hypothesized to be similar to one another but dissimilar from the remaining positions (i.e., positions 3 and 4); **D**, Within-modality • Boundary-position model, in which the patterns between modalities are hypothesized to be dissimilar, and boundary positions (i.e., positions 1 and 4) are hypothesized to be similar to one another but dissimilar from the remaining positions (i.e., positions 2 and 3).

Regarding the modality factor, based on the hypothesis that spatialization is not modality-specific, our favored hypothesis was that the activity patterns would be similar across the visual and auditory conditions, rather than dissimilar. We also hypothesized that distinct patterns would emerge between the early and late positions of a sequence, rather than between middle and boundary positions. This hypothesis corresponds to the mental whiteboard hypothesis (Abrahamse et al., 2014), suggesting that position marking is achieved using spatial coordinates within an internal space. Different activation patterns should then be detected between the first positions in the sequence (probe position 1 and 2) and the last ones (probe position 3 and 4), because coordinates for consecutive positions should be more similar than coordinates for distant positions^3^. This prediction is also congruent with several fMRI findings in the domain (e.g., Attout et al., 2022; Cristoforetti et al., 2022). We thus hypothesized that the *Across-modality • Early-late position* model (model A in Figure 2) would offer the best fit for the neural data.

By including models based on different theoretical perspectives (i.e., models B, C, and D), we aimed to test whether the observed similarity between the neural and theoretical RDMs reflects a reliable cognitive structure rather than coincidental correlations. As models based on different theoretical assumptions can nonetheless show significant associations with neural data, it is crucial to test multiple models and ensure that their fit differ significantly (Nili et al., 2014). Specifically, we aimed to ensure that similarity with the favored model (*Across-modality • Early-late position* model) significantly exceeded that with a control model, thus ensuring that the favored model provides a specific account of the representational pattern.

Rank-based correlation tests were performed to assess the relationship between the theoretical models and the individual neural RDMs within each ROI. This analysis was performed by comparing the lower triangular portions of the neural and theoretical dissimilarity matrices. To assess whether each theoretical model significantly predicted the neural RDMs in each ROI, we conducted a linear mixed-model analysis (Dimsdale-Zucker & Ranganath, 2018; Nili et al., 2014). In this model, the Fisher *z*-transformed Spearman correlation values were entered as predicted variables, with the theoretical models, ROIs, and their interaction as predictors, and participant as a random effect. Post-hoc tests were then conducted to assess the significance of each model within each ROIs, and subsequently pairwise comparisons between models were carried out to determine whether one theoretical model predicted the neural data significantly better than the others. For these post-hoc tests, the significance threshold was set at *p* < .05, corrected for multiple comparisons using false discovery rate (FDR) correction. This analysis was performed separately for the recognition and encoding phases.

## Results

### Behavioral results

The data from one participant was discarded in the auditory condition due to outlier performance (68% accuracy), resulting in a total of 40 participants in the auditory condition and 41 participants in the visual condition. The mean response time and proportion of hits per condition is reported in Table 1. Participants tended to respond faster in the visual condition than in the auditory condition (*t*(39) = 6.96, *p* < .001, Cohen’s *d* = 1.1), while their accuracy was similar (*t*(39) = −1.53, *p* = .133, Cohen’s *d* = 0.24).

**Table 1.** Percentage of correct responses (with standard deviation) and mean response time (in milliseconds, with standard deviation) for each experimental condition

|  | Proportion of<br>correct responses (SD) | Response times (SD) |
| --- | --- | --- |
| Auditory Condition | 93% (5%) | 1213 ms (245 ms) |
| Visual Condition | 94% (4%) | 1035 ms (228 ms) |

To investigate the presence of a spatialization effect, an ANOVA was first conducted. As the distribution of response times was positively skewed (*skewness* = 1.52), we used a log-transformation of the response times as dependent variable (see Guida et al., 2020; Oberauer, 2005). The factors included in this ANOVA were the hand of response, position of the item in the sequence, and condition. The results indicated a main effect of condition (*F* (1, 39) = 60.01, *p* < .001, *η* = 0.61), further confirming that participants responded faster in the visual than auditory modality (see Table 1). A main effect of the response hand (*F* (1, 39) = 9.44, *p* = .004, *η* = 0.19) and of the position of the item in the sequence (*F* (2.56, 99.73) = 13.02, *p* < .001, *η* = 0.13) were also observed. Participants tended to respond faster with their right hand than with their left hand (*M* = 1081 *ms*, *SD* = 203 *ms*, against *M* = 1115 *ms*, *SD* = 224 *ms*, respectively). In addition, participants responded faster to the first and last positions than to the middle positions (*M* = 1078 *ms*, *SD* = 206 *ms* and *M* = 1055 *ms*, *SD* = 202 *ms* for positions 1 and 4, respectively; against *M* = 1150 *ms*, *SD* = 242 *ms* and *M* = 1112 *ms*, *SD* = 227 *ms* for positions 3 and 4, respectively). Details of the results of this ANOVA are presented in Table S2 in Supplementary materials.

Crucially, results also showed a significant interaction between hand and position (*F* (2.52, 98.31) = 4.62, *p* = .007, *η* = 0.11), but only a marginally significant interaction with the condition (*F* (2.77, 108.11) = 2.61, *p* = .06, *η* = 0.06). These results suggest the presence of a spatialization effect, which was not modulated by the condition. To further investigate the presence of spatialization in each condition, linear regressions were conducted on the difference of mean response times between the time taken to respond using the right hand minus the time taken to respond using the left hand, as a function of the position of the item in the sequence (see e.g., van Dijck and Fias, 2011). These regressions highlighted a significant effect in the visual condition (*t*(160) = −3.11, *p* = .002, *BF*_10_ = 14.6, with a mean decrease of −35.1 ms per position), whereas the effect was not significant in the auditory condition (*t*(155) = −0.88, *p* = .381, *BF*_01_ = 4, with a mean decrease of −9.5 ms per position), as shown in Figure 3. However, the absence of effect in the auditory condition is difficult to interpret as the background noise induced by the MRI scanner could have influenced the results, in particular because spatialization has previously been detected using auditory stimuli (Bottini et al., 2016; Guida & Porret, 2022; Guida et al., 2016).

**Figure 3.**
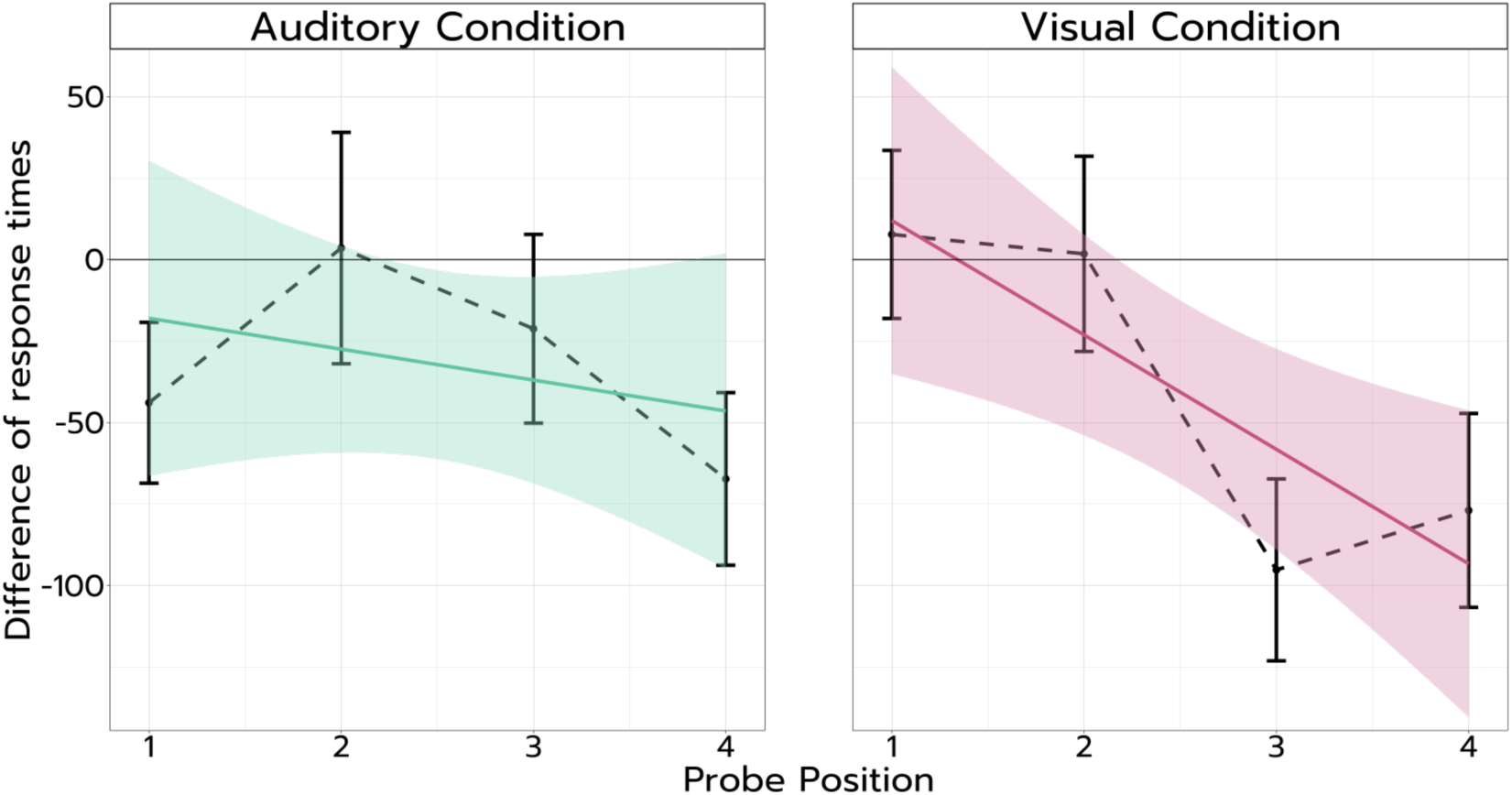
Difference of response times between right and left hands as a function of the position of the item in the sequence, and associated linear regression for each experimental condition. Note. The figure has been produced using aggregated data: for each participant, an average d_RT_ per position was obtained by calculating the difference between the mean_RT_ obtained with the right hand and the mean_RT_ obtained with the left hand. The error bars correspond to ±1 standard error. d_RT_ are expressed in milliseconds.

### Neuro-imaging results: univariate analyses

#### Whole-brain analyses

We first conducted whole-brain analyses to identify the regions associated with spatialization within each modality, and to assess potential differences across conditions through direct comparisons between our two modalities. We assessed the overall activation during the encoding and recognition phases across both conditions by computing one sample *t*-tests at whole-brain level. The results are reported in Table 2 and shown in Figure 4 (panels A and B). During the encoding phase, the visual condition elicited activation in the left anterior insula, left postcentral gyrus, as well as in the right inferior occipital gyrus, right transverse temporal gyrus (Heschl’s gyrus) and in the cerbellum. The auditory condition elicited activation in the left inferior frontal gyrus, left SMA, left IPS, and in the right precentral gyrus. Overall, these results thus show a fronto-parietal network typically associated with working memory processing, as well as activations in item-processing areas such as the occipital gyri and Heschl’s gyri, congruent with previous studies.

**Figure 4.**
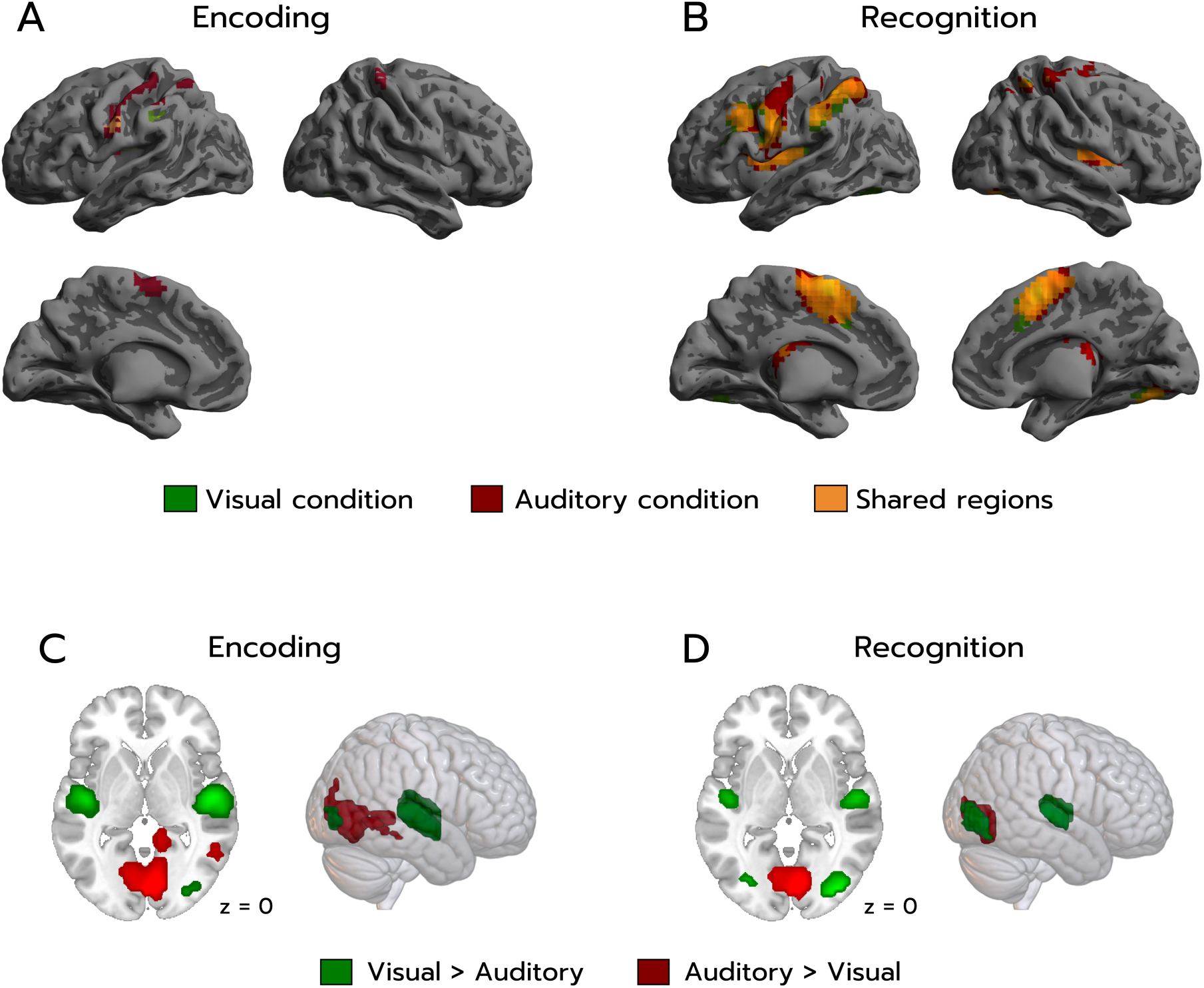
Whole-brain activation maps for the encoding and recognition phases. Panels **A** and **B** display regions activated by i) in green, the visual condition versus the implicit baseline, ii) in red, the auditory condition versus the implicit baseline, and iii) in orange, the conjunction of both conditions versus the implicit baseline, for the encoding and recognition phases, respectively. Panels **C** and **D** show cerebral regions whose activity was elicited by the contrast i) in green, [VISU > AUDIO], and ii) in red, [AUDIO > VISU], also for the encoding and recognition phases, respectively. Note. Neural activity is projected onto 2D slices and 3D inflated anatomical templates using the MNI2FS toolbox (Price, 2024), using a threshold of p< .05 (FWE-corrected), with a minimum cluster size of k = 10 voxels.

**Table 2.**
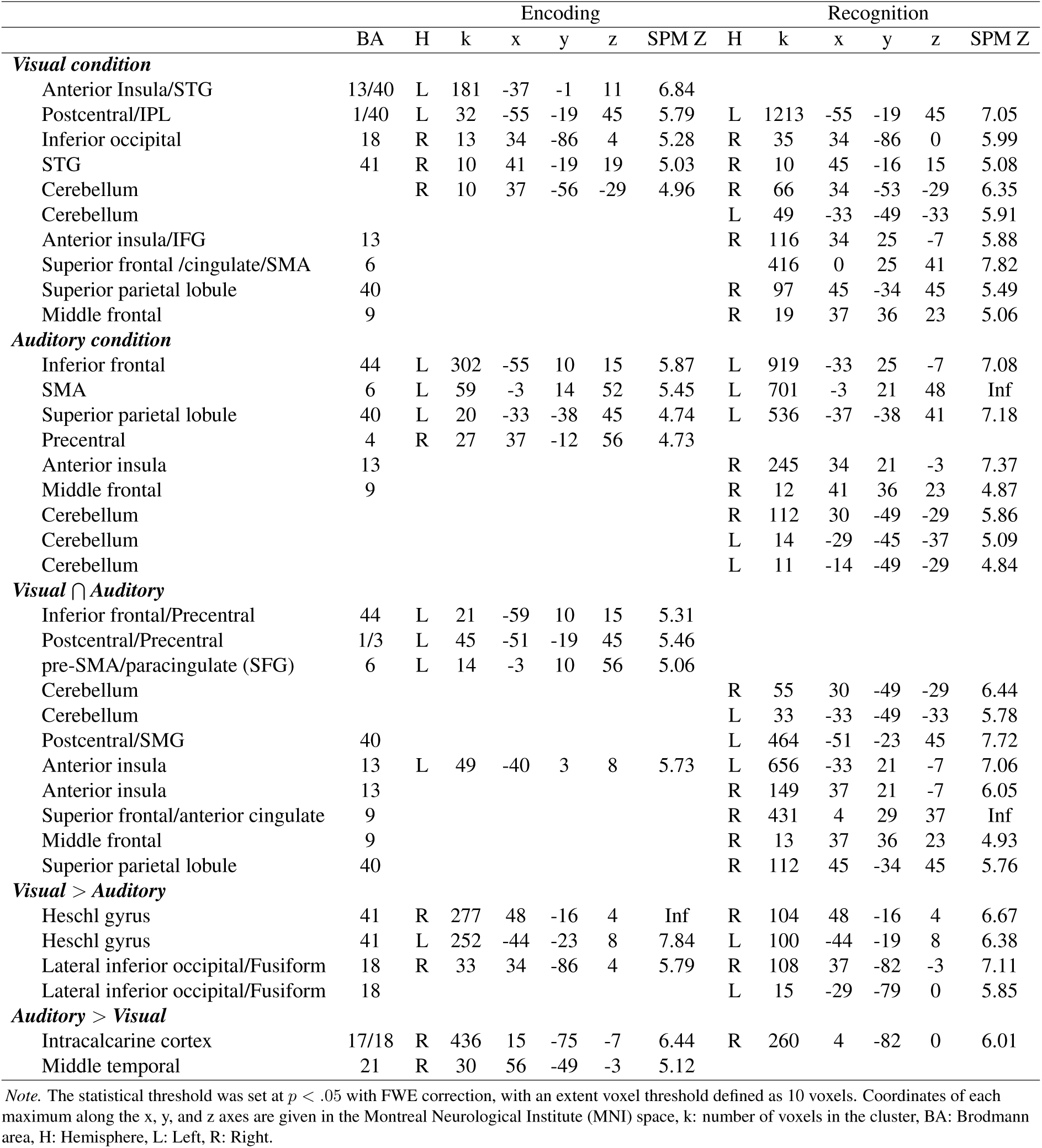
Significant activation foci for the encoding and recognition phases in the visual and auditory spatialization tasks, conjunction analysis (common activation across both tasks), as well as contrasts between tasks

During the recognition phase, activation was observed for the visual condition in the right inferior occipital gyrus, right anterior insula, right middle frontal gyrus, and in the right IPS. Activation was also observed bilaterally in the SMA and cerebellum, in the left postcentral gyrus and transverse temporal gyrus (Heschl’s gyrus). For the auditory condition, activation was noted in the right anterior insula, middle frontal gyrus, and in the left IPS and inferior frontal gyrus. In addition, activation was observed bilaterally in the SMA and cerebellum. Again, this network is largely consistent with the network found in previous working memory studies.

We then conducted a conjunction analysis to investigate the regions showing shared activation across both conditions for each phase (see Table 2). The conjunction analysis revealed shared activation during the encoding phase in frontal and inferior parietal areas comprising the anterior insula, inferior frontal gyrus, pre-central and post-central gyri, as well as the pre-SMA and paracingulate gyrus. Across both modalities, the activation in the inferior frontal gyrus (BA44, Broca’s area) and in the premotor cortex and SMA (BA6) could correspond to the shared processes comprising phonological coding and rehearsal of the sequence (Henson et al., 2000). During the recognition phase, shared activation was observed in the left supramarginal gyrus, right superior and middle frontal gyri, right superior parietal lobule, as well as bilaterally in the cerebellum and anterior insula. These analyses thus revealed a similar activation pattern between conditions and phases of the task, although the network was larger during recognition than encoding. The regions activated were consistent with the fronto-parieto-cerebellar network observed in previous research (e.g., Attout et al., 2014; Henson et al., 2000; Majerus et al., 2010; Marshuetz et al., 2000). This result also tends to indicate that the way in which information is processed in the SPoARC cannot be differentiated based on the presentation modality. These activations are shown in Figure 4 (panels A and B).

Finally, direct comparisons were computed between the two conditions to identify potential modality-specific activations across the whole brain. The contrast [VISU > AUDIO] showed increased activity in Heschl’s gyrus and in the right lateral occipital gyrus in both the encoding and recognition phases. Activation in Heschl’s gyrus was mostly attributable to a reduced activation in the auditory condition compared to the baseline, potentially reflecting suppression of the background noise of the scanner to facilitate higher-order speech processing. Activation in the lateral inferior occipital gyrus included the fusiform gyrus, suggesting both higher-order processing and involvement of visual memory. In addition, when performing the opposite contrast ([AUDIO > VISU]), we found significant activation in the intracalcarine cortex for both phases, and in the middle temporal gyrus during the encoding phase. Activation in the intracalcarine cortex was however mostly due to a reduced activation in the visual condition compared to the baseline, potentially reflecting a greater focus on the central fixation point (where stimuli were presented) and suppression of peripheral visual input (see retinotopic organization in the occipital cortex, Wandell et al., 2007). These effects are also reported in Table 2, and shown in Figure 4 (panels C and D).

#### ROI analyses

Next, ROI analyses were carried out to directly compare the overall response magnitude across voxels within working memory regions, in order to assess whether spatialization engages the same neural mechanisms across modalities. We thus applied the contrasts between conditions within our predefined ROIs, revealing an increased activation in the posterior IPS and in the IFG in the auditory condition during the encoding phase (*t*(40) = 3.49, *p* = .001, and *t*(40) = 3.77, *p* < .001, respectively). Note that although accuracy did not differ between conditions, response times were shorter in the visual modality, potentially indicating lower attentional demands compared to the auditory condition (Majerus, Poncelet, Van der Linden, et al., 2006). This attentional difference might thus partly explain the greater activation observed for auditory encoding. No other significant effects were observed across both the encoding and recognition phases (all *t_s_ <* 2.54, all *p_s_ > .*015 for the encoding phase, and all *t_s_ <* 2.98, all *p_s_ > .*005 for the recognition phase, with the *p*-value threshold set at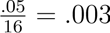). The largely similar activation levels within our ROIs are consistent with the whole-brain results, suggesting minimal differences in the networks typically associated with working memory processing across conditions. Although significant differences were observed in two ROIs during the encoding phase, these effects were limited and did not suggest widespread differences in working memory-related networks.

### Neuro-imaging results: multivariate pattern analysis

Subsequently, we conducted multivariate pattern analyses within ROIs, enabling to provide a more nuanced approach by considering patterns of activity across multiple voxels. This method enabled to examine whether these patterns are shared across conditions and to determine whether patterns associated with the encoding or recognition of specific serial positions (early *vs.* late positions in a sequence) could be discriminated, ultimately indicating that the corresponding cortical region carries information regarding the order of elements in a sequence.

By assessing the similarity between our neural and theoretical RDMs during the encoding phase, we found that significant mean similarity was observed between the neural data and all four theoretical models in the superior frontal gyrus (*t*(1279) ≥ 2.32, *p_FDR_ ≤ .*03, *d* ≥ 0.36; see Table S3 in Supplementary materials), with no significant differences between models (*t*(1240) ≤ 1.12, *p_FDR_ ≥ .490, *d* ≥* 0.25; see Table S4 in Supplementary materials). In the middle frontal gyrus, significant mean similarity was also observed between the neural data and all theoretical models except the *Within-modality • Boundary-position* model (*t*(1279) ≥ 2.48, *p_FDR_* ≤ .021, *d* ≥ 0.39; against *t*(1279) = 1.79, *p_FDR_* = .099, *d* = 0.28, respectively). Again, no significant differences between models were observed for this region (*t*(1240) ≤ 2.26, *p_FDR_* ≥ .223, *d* ≤ 0.50). For the inferior frontal gyrus, significant mean similarity was observed between the neural data and all theoretical models except with the *Across-modality • Boundary-position* model (*t*(1279) ≥ 4.54, *p_FDR_* < .001, *d* ≥ 0.69; against *t*(1279) = −1.16, *p_FDR_* = .291, *d* = 0.18, respectively). Pairwise comparisons revealed that the *Within-modality • Early-late position* model showed a significantly higher mean similarity than all other models in this region (*t*(1240) ≥ 3.24, *p_FDR_* ≤ .003, *d* ≥ 0.72). Mean correlations between neural and theoretical RDMs across ROIs are presented in Figure 5. The findings from the current set of ROIs suggest that the inferior frontal gyrus shows differences in patterns of activity both across conditions and between early and late item positions. In contrast, no conclusive evidence was found for pattern differences in the middle and superior frontal gyri across either conditions or sequence positions.

**Figure 5.**
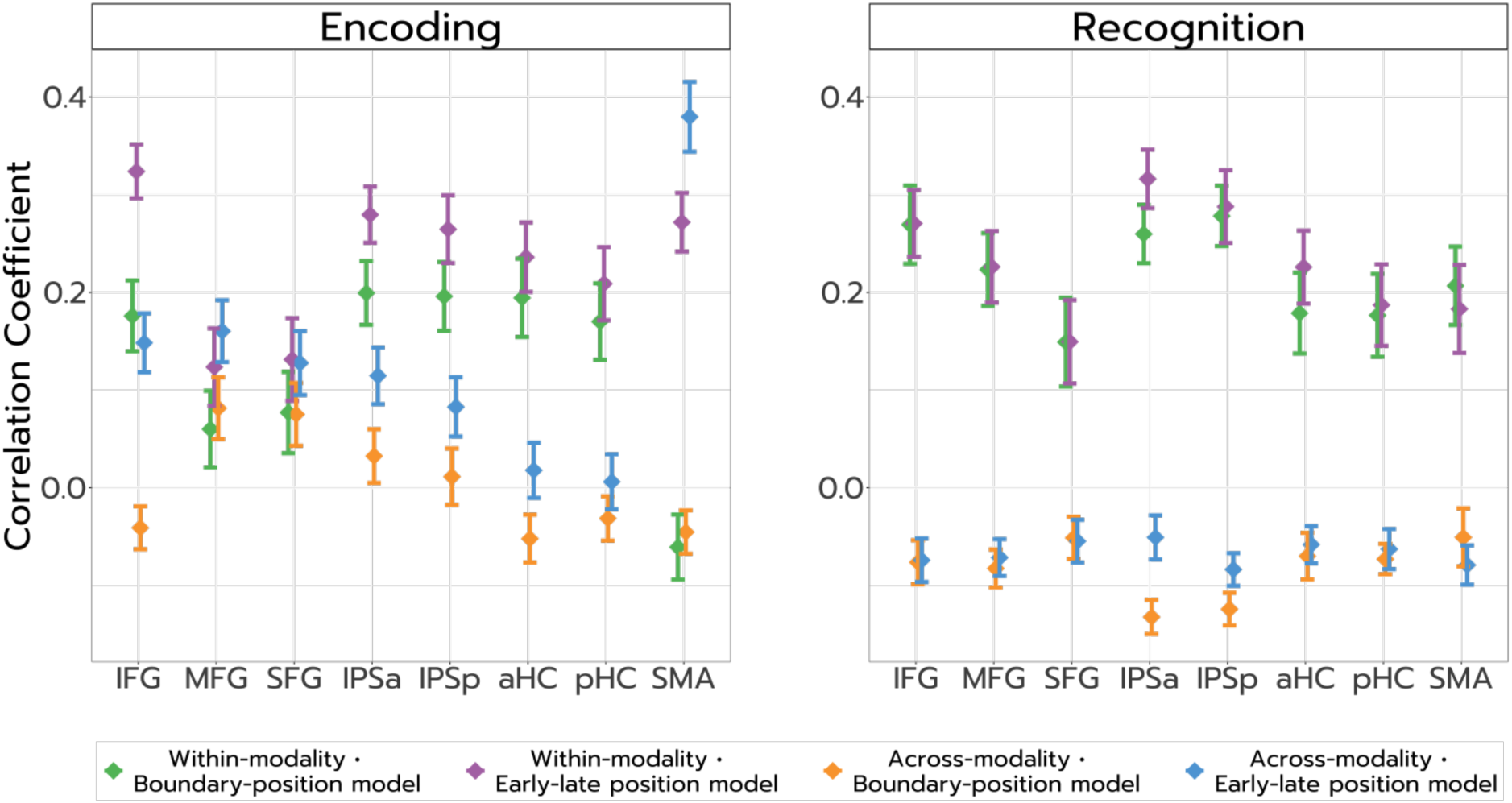
Spearman correlation coefficients between the observed RDMs and the four theoretical RDMs across regions of interest, for the encoding and recognition phases. Note. The error bars correspond to ±1 standard error.

In the parietal ROIs, we observed a significant similarity between neural patterns and the *Within-modality • Early-late position* model (*t*(1279) ≥ 8.00, *p_FDR_ < .*001, *d* ≥ 1.12). Significant, although smaller, similarities were also found for the *Within-modality • Boundary position* and *Across-modality • Early-late position* models (*t*(1279) ≥ 2.48, *p_FDR_* ≤ .021, *d* ≥ 0.39). Pairwise model comparisons revealed that the *Within-modality • Early-late position* and *Within-modality • Boundary position* models did not significantly differ in mean similarity (*t*(1240) ≤ 1.76, *p_FDR_* ≥ .153, *d* ≤ 0.39). However, in the posterior IPS, both of these models showed greater similarity with neural patterns than the two remaining models (*t*(1240) ≥ 2.41, *p_FDR_* ≤ .046, *d* ≥ 0.53). In contrast, in the anterior IPS, only the *Within-modality • Early-late position* model demonstrated greater similarity than the other two models (*t*(1240) ≥ 3.55, *p_FDR_* ≤ .002, *d* ≥ 0.79). These RSA results indicate condition-related differences represented in parietal regions, whereas no clear conclusions could be drawn regarding the position of the item in the sequence.

Next, we examined the results in the hippocampal ROIs. A significant similarity was observed between the observed neural patterns and both the *Within-modality • Early-late position* and *Within-modality • Boundary position* models (*t*(1240) ≥ 5.09, *p_FDR_ < .*001, *d* ≥ 0.80). These two models could not be statistically distinguished from one another (*t*(1240) ≤ 0.87, *p_FDR_* ≥ .490, *d* ≤x 0.19), but each showed significantly greater similarity with the neural RDMs than the two remaining models (*t*(1240) ≥ 3.37, *p_FDR_* ≤ .003, *d* ≥ 0.74). These results suggest that the observed similarity in the hippocampus is primarily driven by condition-related differences, with no conclusive evidence for position-based coding. In the supplementary motor area (SMA), both the *Across-modality • Early-late position* and *Within-modality • Early-late position* models showed strong similarity with neural patterns (*t*(1240) ≥ 8.23, *p_FDR_ < .*001, *d* ≥ 1.29). However, the *Across-modality • Early-late position* model exhibited higher similarity with the neural data than model *Within-modality • Early-late position* (*t*(1240) = 2.96, *p_FDR_* = .007, *d* = 0.65). These findings highlight the dominant role of sequence (temporal) processing in the SMA, supporting distinct neural patterns for the beginning versus the end of a sequence, while also suggesting a tendency to encode position information in a modality-independent manner.

Given that we observed distinct patterns of activity between conditions in most ROIs, an exploratory analysis per modality was conducted to assess potential modality-specific position effects. For this analysis, the data were restricted to the four runs corresponding to a single condition, resulting in two 4 × 4 matrix per ROI for each participant (i.e., a 4 positions × 4 positions matrix for both the visual and auditory modalities). These neural RDMs were then compared to both an *Early-late position* model and a *Boundary position* model. Again, the similarity between each neural and theoretical RDM was assessed using Spearman rank correlations, which were subsequently *z*-transformed. These values were entered as the dependent variable in linear mixed models, with the model, ROI, and their interaction as fixed effects, and participant included as a random effect. Separate models were fit for each condition.

In the auditory condition, significant similarity with the *Early-late position* model was observed in the inferior frontal gyrus (IFG), superior frontal gyrus (SFG), and posterior hippocampus (pHC), although these effects reached significance only when considering uncorrected *p*-values (*t*(621) ≥ 2.02, *p_uncorr._* ≤ .044, *d* ≥ 0.32; see Table S5 in Supplementary materials for details of the results). Moreover, the *Early-late position* model had a significantly higher similarity with the neural data than the *Boundary position* model in the IFG and pHC (*t*(600) ≥ 2.60, *p_FDR_* ≤ .020, *d* ≥ 0.58), as well as in the anterior hippocampus (aHC; *t*(600) = 2.58, *p_FDR_* = .020, *d* = 0.57; see Table S6 in Supplementary materials). These findings suggest that, in the auditory condition, activity patterns in the IFG and pHC (and to a lesser extent in the SFG and aHC) differentiated between the early and late positions in a sequence. In the visual condition, significant similarity with the *Early-late position* model was found in the IFG, middle frontal gyrus (MFG), and both anterior and posterior parts of the intraparietal sulcus (IPSa and IPSp; *t*(640) ≥ 2.81, *p_FDR_* ≤ .014, *d* ≥ 0.44). This model also showed a higher similarity with the neural data than the *Boundary position* model in the IFG and IPSa (*t*(600) ≥ 2.36, *p_FDR_* ≤ .049, *d* ≥ 0.52), and in the IPSp when considering uncorrected *p*-values (*t*(600) = 2.24, *p_uncorr._* = .026, *d* = 0.49). Finally, across both sensory conditions, neural patterns in the supplementary motor area (SMA) showed robust similarity with the *Early-late position* model (*t*(621) ≥ 6.42, *p_FDR_ < .*001, *d* ≥ 1.03). This model also showed a higher mean similarity with the neural data than the *Boundary position* model (*t*(600) ≥ 6.86, *p_FDR_ < .*001, *d* ≥ 1.51), consistent with the analysis ran across both conditions. These results, presented in Figure 6, suggest distinct patterns of neural activity between the beginning and the end of a sequence in the IFG and IPSa for the visual condition, and in the IFG and pHC for the auditory condition.

**Figure 6.**
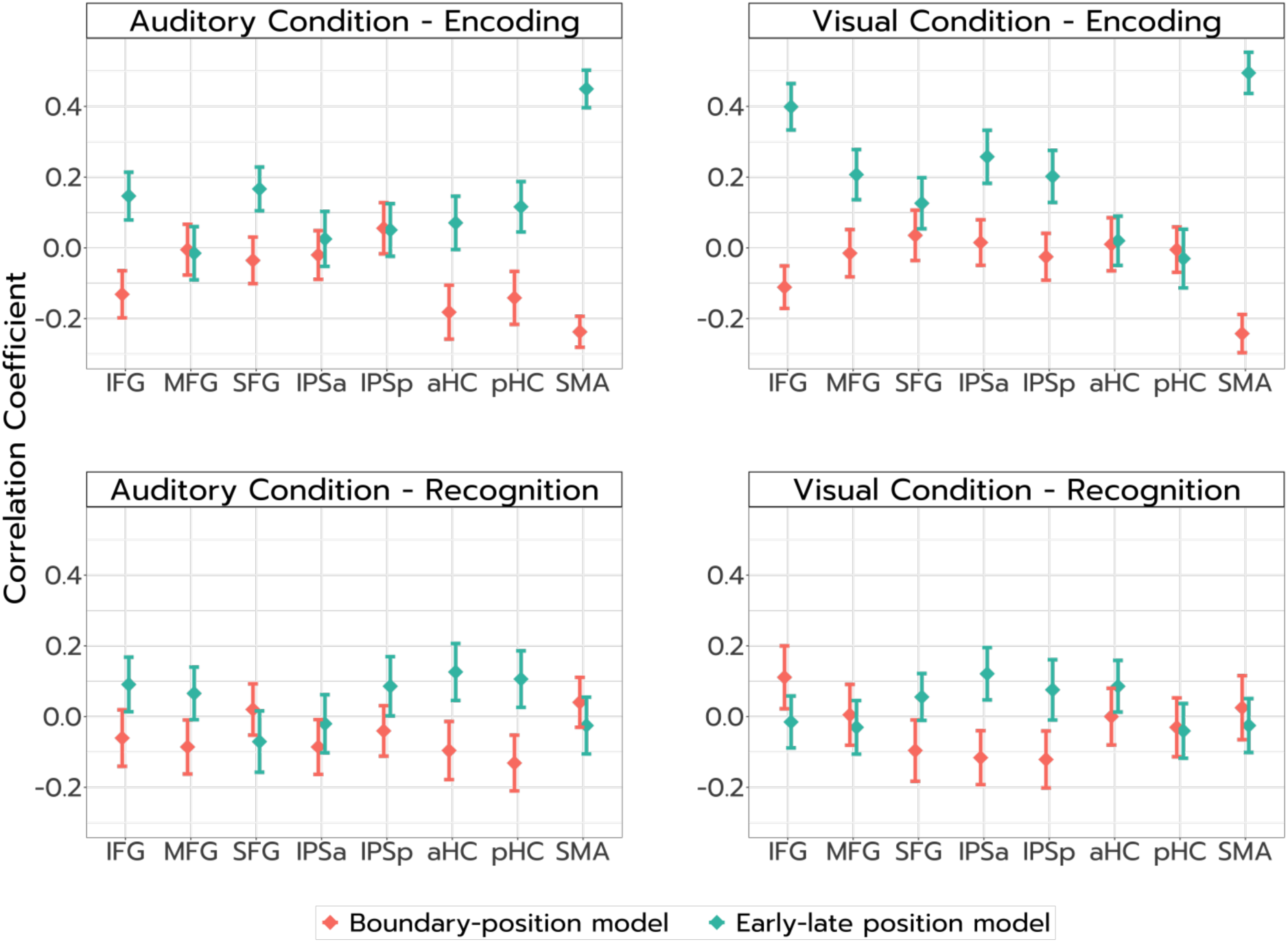
Spearman correlation coefficients between the observed RDMs and the two theoretical RDMs across regions of interest, for the encoding and recognition phases and each condition separately. Note. The error bars correspond to ±1 standard error.

Overall, our results regarding the encoding phase revealed that, although the visual and auditory conditions showed different patterns of activation in the inferior frontal gyrus, they both elicited different patterns between early and late positions in the sequence. In addition, different patterns were observed between conditions in the intraparietal sulcus and the hippocampus. Specifically, distinct patterns for early versus late items were observed in the anterior IPS for the visual condition and in the posterior hippocampus for the auditory condition.

Subsequently, we conducted the same analysis for the recognition phase. We found that both the *Within-modality • Early-late position* and *Within-modality • Boundary position* models showed a significant correlation with the neural data across all ROIs (*t*(1273) ≥ 4.85, *p_FDR_ < .*001, *d* ≥ 0.16, see Table S3). However, there was no significant difference in mean similarity between the two models (*t*(1240) ≤ 1.40, *p_FDR_* ≥ .229, *d* ≤ 0.31). Both models showed higher similarity with the neural data compared to the other two models (*Across-modality • Early-late position* and *Across-modality • Boundary position*, *t*(1240) ≥ 4.57, *p_FDR_ < .*001, *d* ≥ 1.01). Given that the results of this analysis appeared to be driven by different patterns of activity across conditions, we also performed the exploratory analysis for each condition separately. However, we found that neither the *Early-late position* nor the *Boundary position* models significantly related to the neural data across all conditions and ROIs (*t*(640) ≤ 1.80, *p_FDR_* ≥ .604, *d* ≤ 0.28, see Table S5). Still, we observed a significant higher similarity with the *Early-late position* model in the anterior and posterior hippocampus for the auditory condition (*t*(600) ≥ 2.03, *p_uncorr._* ≤ .043, *d* ≥ 0.45), and in the anterior IPS for the visual condition (*t*(600) = 2.26, *p_uncorr._* = .024, *d* = 0.50), when considering uncorrected *p*-values. The results of these analyses are shown in Figure 5 (combining the effects of condition and position) and Figure 6 (for the effect of position in each condition separately).

Overall, these findings suggest that during recognition, both conditions elicited distinct patterns of activation. Although the results regarding the effect of serial position were more contrasted, potentially due to the more restricted dataset for this phase, there was a tendency for different patterns between the beginning and end items of a sequence in the anterior IPS for the visual condition, and in the anterior and posterior hippocampus for the auditory condition, similar to the findings for the encoding phase.

## Discussion

The objectives of the current study were two-fold. First, we aimed to show that the network involved in the processing of order information in the SPoARC is modality-independent, while also showing distinct networks related to the processing of item information in sensory areas. Second, we intended to investigate how order is coded across modality by investigating whether frontal, parietal, as well as hippocampal regions show sensibility to different serial positions in both the visual and auditory modalities. To investigate this, we performed a direct comparison between an auditory and visual SPoARC task in an fMRI setting. Using a representational dissimilarity analysis, we hypothesized that activation patterns would be similar across modalities within working memory-related ROIs. This representational dissimilarity analysis also investigated the effect of the position of the item in the sequence. In line with previous findings, we expected to observe different patterns of activation between early and late working memory positions within the fronto-parietal network and the hippocampus (Cristoforetti et al., 2022).

First, at the behavioral level, we observed a significant SPoARC effect in the visual condition but not in the auditory condition. The absence of effect in the auditory condition is difficult to interpret, as background noise from the MRI scanner could have interfered with stimulus processing. Indeed, this absence of effect could be due to two distinct factors. One possibility is that the acoustic environment of the scanner increased task difficulty, as indicated by the longer response times in the auditory compared to the visual condition (although we still observed similar performance in terms of accuracy), thus preventing the measure of more spontaneous response times necessary to detect the effect. This interpretation is also consistent with previous studies detecting the SPoARC with auditory stimuli when outside the scanner (Bottini et al., 2016; Guida & Porret, 2022; Guida et al., 2016). Alternatively, the difference between the visual and auditory conditions might reflect distinct memorization processes across sensory modalities, as suggested by the different patterns of activations identified using RSA. The exact mechanisms underlying these modality-specific processes are developed later, in light of the multivariate findings below.

Second, despite the absence of a SPoARC effect in the auditory condition, the results of the univariate analysis showed that both modalities elicited activation within a fronto-parieto-cerebellar network corresponding to the previously identified working memory circuits (Chai et al., 2018; Cowan et al., 2011; Cristoforetti et al., 2022; Kalm & Norris, 2014; Majerus et al., 2018). These findings thus support the existence of a shared cortical network for processing ordinal information across sensory modalities. In addition to this shared cortical network supporting domain-general functions, such as order processing or attentional control, we also observed specific differences of activation in early sensory cortices across conditions, suggesting differential processing of item information within sensory regions (Albers et al., 2013; Chai et al., 2018; Cowan et al., 2011; Majerus et al., 2010; Riggall & Postle, 2012; Vetter et al., 2014). This interpretation is further supported by studies showing successful decoding of stimulus identity in sensory cortices (visual or auditory), while decoding in frontal, parietal, or motor regions was unsuccessful (Albers et al., 2013; Riggall & Postle, 2012; Vetter et al., 2014). This dissociation between sensory and fronto-parietal regions also aligns with positional models of working memory, which propose distinct codes for item and order information (Brown et al., 2000; Henson, 1998; Oberauer, 2009). Overall, the univariate findings reinforce the view that order processing in working memory relies on a shared fronto-parieto-cerebellar network across modalities, whereas item processing relies on modality-specific sensory regions, and further extend this framework to a SPoARC task.

Third, beyond these univariate findings, our representational similarity analysis allowed us to identify modality-specific patterns of activation, and, importantly, to reveal for the first time distinct patterns of activation for early versus late items in different regions depending on sensory modality. Such results suggest that order processing would rely on partially distinct mechanisms across modalities rather than on shared processes. Indeed, while activation levels were similar across conditions within our regions of interest, a multivariate approach revealed distinct neural patterns for visual and auditory information in fronto-parietal regions and the hippocampus, suggesting modality-specific representations within largely overlapping networks. In particular, we identified distinct patterns in the inferior frontal gyrus, in both the anterior and posterior parts of the intraparietal sulcus and of the hippocampus during encoding, as well as in all frontal regions (inferior, middle, and superior frontal gyri), and in both the anterior and posterior parts of the intraparietal sulcus and of the hippocampus during recognition. Our results contradict those of Rizza et al. (2024), who found no significant pattern differences in the fronto-parietal network when comparing the maintenance of auditory versus visual-spatial information. Still, it could be argued that their use of non-verbal stimuli and the shared spatial component of the task across modalities might have contributed to the observed similarity (see Hurlstone, 2019, for differences in position coding across verbal and spatial sequences). Moreover, their multivariate analyses focused only on the maintenance phase, which we did not model here. In contrast, our results are consistent with a study by Majerus et al. (2018), in which attentional control and encoding load could not be reliably decoded across a visual and an auditory presentation (by training a classifier on one modality and testing it on the other). In addition, although contrary to our hypothesis, this finding can be related to studies supporting the shared involvement of frontal and parietal regions for order processing across domains, while also pointing to domain-specific activation patterns within these regions. For example, regions such as the IPS and frontal gyri have been shown to be consistently involved in ordinal processing across stimuli including digits, letters, shapes, and working memory sequences (Attout et al., 2014; Fias et al., 2007; Fulbright et al., 2003). These regions are also activated during the processing of long-term memory knowledge, including the magnitude and order of digits (Fias et al., 2003; Franklin & Jonides, 2009; Sokolowski et al., 2017). However, a re-analysis of the results of Fias et al. (2007) by Zorzi et al. (2011) revealed distinct activation patterns for ordinal judgments involving letters versus digits. Similarly, Attout et al. (2022) reported different patterns between ordinal processing of digits, letters, and working memory sequences (although no distinction was found between letters and working memory sequences). Overall, the results of the MVPA analysis, showing similar levels of activations across modalities while still showing differences in neural patterns, suggest that despite the modality-general involvement of working memory regions (consistent with prior studies Arnott et al., 2005; Kalm and Norris, 2014; Majerus et al., 2018; Zhang et al., 2003), order information within the working memory network still carries information regarding the input format of information to memorize.

In addition to the condition-related pattern differences, we also observed distinct activation patterns based on the position of items in working memory. These patterns were analyzed separately for each modality, given the observed differences in neural pattern between conditions. First, during the encoding phase, we found position-related pattern differences in both the inferior frontal gyrus and the supplementary motor area, suggesting different patterns between early and late positions in working memory. This result was consistent across conditions. Additionally, modality-specific effects emerged: in the visual condition, different activation patterns were observed between early and late positions in the anterior intraparietal sulcus. In contrast, in the auditory condition, these effects were localized to the posterior hippocampus. No significant position-related effects were found in the other frontal regions (middle frontal gyrus or superior frontal gyrus). A similar pattern was observed during the recognition phase, although the results were non-significant after FDR correction. Specifically, we found a trend toward discriminable patterns for early versus late positions in the anterior IPS in the visual condition, and in the hippocampus (both anterior and posterior parts) in the auditory condition.

Based on a framework put forward by Bottini and Doeller (2020), we thus propose that the representation of order preferentially relies on distinct cortical regions depending on the modality of presentation. In their review, Bottini and Doeller (2020) suggested that conceptual knowledge could be represented spatially by using self-centered image spaces (i.e., an egocentric reference frame) or world-centered cognitive maps (i.e., an allocentric reference frame). These self-centered image spaces, supported by the parietal cortex (Bottini & Doeller, 2020), have only one or two dimensions and can, for example, take the form of a left–right or top–down axis, as seen in representations such as the mental timeline or mental number line (Boroditsky, 2000; Galton, 1880; Weger & Pratt, 2008). In contrast, the world-centered cognitive maps correspond to more complex representations of conceptual knowledge (i.e., internal spatial representations organized in an allocentric reference frame) and are thought to rely on the activity of the hippocampus (Bellmund et al., 2018; Bottini & Doeller, 2020), which is primarily associated with spatial navigation (Bicanski & Burgess, 2018; Maguire et al., 1998), but has also been shown to encode non-spatial conceptual dimensions (e.g., sound frequency, Aronov et al., 2017). We thus suggest that the intraparietal sulcus, in relation with the use of low-dimensional spaces, would be more engaged with visual presentations, whereas the hippocampus, in relation with the use of high-dimensional spaces, would be more engaged with auditory presentations.

This hypothesis of differential parietal and hippocampal involvement depending on the presentation modality is consistent with previous studies showing decoding of order-related information in the IPS when using visual presentations (e.g., Attout et al., 2022; Cristoforetti et al., 2022). In particular, to the best of our knowledge, only one study has directly examined item position within a sequence using an MVPA analysis (in a visual detection task), showing distinct activation patterns for the first versus last position in a sequence in the left IPS (Cristoforetti et al., 2022). However, the authors were also able to decode the first versus last position of a sequence in the hippocampus. Thus, it is likely that both the IPS and hippocampus contribute to the encoding of item position across sensory modalities (see also Dolfen et al., 2024, suggesting that the hippocampus encodes the temporal order of items regardless of modality). Still, this does not preclude the possibility that certain regions are preferentially engaged depending on the modality.

We thus postulate that order coding within the IPS could be related to the concept of self-centered image spaces (Bottini & Doeller, 2020). As these self-centered images only have one or two dimensions, they could be used to code order along a mental horizontal line (i.e., a one-dimensional left-right axis, or an axis within a 2D space). We additionally propose that order decoding in the IPS might reflect the use of spatial codes for order as described by the mental whiteboard hypothesis (Abrahamse et al., 2014). The mental whiteboard hypothesis proposes that positional markers would be spatial in nature (Abrahamse et al., 2014). Order coding would be achieved by using spatial tags within an internal spatial frame (i.e., the mental whiteboard, which would correspond to a self-centered image space); and spatial attention would be used to retrieve these codes and their associated items. The IPS would thus serve as a key region supporting spatial coding of order, as it is thought to support both order coding (in its anterior portion) and spatial attention (in its posterior portion, which is part of the dorsal attention network). Although spatial attention processing in this area is thought to primarily relate to external spatial attention (Gillebert et al., 2011; Vandenberghe et al., 2001; Yantis et al., 2002), Rasoulzadeh et al. (2021) showed that the same frontoparietal regions are recruited when engaging spatial attention in both external and internal space, as well as for visuo-spatial working memory^4^ (see also Nobre et al., 2004). Our proposition is also consistent with previous studies suggesting that the IPS functions as a spatial template for representing order or directing attention to spatially encoded positions (Attout et al., 2022; Cristoforetti et al., 2022; Zhou et al., 2021). Furthermore, under the hypothesis that order decoding in the IPS reflects the use of these low-dimensional image spaces, we argue that the preferential involvement of this region for visually presented information may result from the physical layout of items on a two-dimensional screen, which would more naturally encourage low-dimensional spatial encoding strategies.

Contrary to the low-dimensional spaces discussed above, we propose that order decoding in the hippocampus would reflect the use of world-centered cognitive maps to represent order information. The hypothesis that order decoding in the hippocampus reflects the use of allocentric spatial codes also aligns with results by Zhou et al. (2021), who reported connectivity between the IPS, hippocampus, and retrosplenial cortex in their fMRI study of the SPoARC^5^. The authors interpreted this network as supporting allocentric representations of order via the hippocampus, with the retrosplenial cortex acting as a functional bridge between allocentric spatial codes and the egocentric reference frame required for manual responses (i.e., responding with the left or right hand). Further extending our hypothesis, we argue that the preferential involvement of this region for auditorily presented information reflects the fact that, as the auditory condition provides no inherent spatial template, it could possibly prompt participants to rely on these more complex allocentric spatial representations to encode the order of items. Alternatively, it is possible that the visual modality encourages the use of spatial codes due to the tangible presentation of items on a screen, whereas the auditory modality, being less spatially grounded, might favor temporal coding strategies, thereby engaging more strongly the left hippocampus (Cristoforetti et al., 2022; Gauthier et al., 2020). Indeed, in the study by Cristoforetti et al. (2022), the authors revealed a decoding of the serial position in the IPS, hippocampus, SMA, and SMG; and suggested that the IPS could be associated with spatial-attentional coding, while the hippocampus, SMG, and SMA could be linked to temporal coding mechanisms. Importantly, Gijssels et al. (2019) provided evidence that spatial mappings are more prominent in a visual modality whereas temporal mappings are more prominent in an auditory modality, consistent with our hypothesis of distinct mappings based on stimulus modality. Overall, the involvement of the hippocampus in order coding in working memory could reflect the use of spatial allocentric or temporal order codes, while the involvement of the intraparietal sulcus could rather reflect the use of low-dimensional egocentric spatial codes.

In addition, this interpretation that the modality-dependent difference of activation patterns across positions reflects the use of distinct representational spaces is also consistent with our behavioral results. Specifically, we found that the SPoARC was detected in the visual but not auditory modality. Recall that we previously suggested that the absence of effect in the auditory condition could be due to two factors, the first one being noise in the scanner, and the second one being distinct memorization processes across sensory modalities (based on the difference of patterns uncovered by the RSA analysis). Indeed, according to our account, when memorizing visual information, low dimensional spaces are preferentially recruited due to the physical layout of items on a two-dimensional screen. As these low dimensional spaces would correspond to a 2D axis such as a mental horizontal line, they would lead to a stronger SPoARC, since this effect captures as left-to-right organization of information. In contrast, when memorizing auditory stimuli, higher-representational spaces would be preferentially recruited. This would lead to a weaker spatialization, as these more complex representations do not correspond to a unique orientation in a left to right direction, as captured in the SPoARC. The difference in spatialization across our two conditions could thus also reflect preferential reliance on distinct representational spaces depending on the input modality. Altogether, our behavioral findings might therefore result from the synergy between these two factors (i.e., the scanner noise and the distinct memorization processes).

However, it should be noted that there are some limitations to the present experiment. First, although participants’ accuracy did not differ across modalities, response times were still faster in the visual condition. This result potentially reflects increased attentional demands when performing the task with auditory material, possibly due to the presence of scanner noise. These increased attentional demands tend to be confirmed by the greater activation in the inferior frontal gyrus and posterior IPS in the auditory condition. This difference in attentional engagement might thus have influenced the multivariate analyses for these specific regions. While dissimilarity analyses using correlation-based distance metrics are largely insensitive to overall activation amplitude (Davis et al., 2014; Oosterhof et al., 2016), differences in attention could still impact the underlying activation patterns across conditions. Further research is thus needed to determine whether the differences in patterns between modalities persist when attention is controlled across conditions. Second, the data available to analyze the effect of serial position in working memory during the recognition phase were inherently more limited than for the encoding phase, which may have reduced the signal-to-noise ratio. This limitation could account for the lack of significant findings regarding the effect of serial position during recognition. Replicating these analyses with a larger dataset would be important to confirm our results. Finally, the SPoARC task we used in this study does not allow for a clear dissociation between item memory and order memory. Investigating this distinction using both visual and auditory modalities could offer valuable insights into the mechanisms underlying order representation in working memory.

## Conclusion

Our results suggest that order coding in working memory preferentially relies on different cortical regions depending on the modality of stimulus presentation. Specifically, positional coding would tend to be preferentially associated with the intraparietal sulcus when memorizing visual materials, whereas it would tend to be preferentially associated with the hippocampus when memorizing auditory materials. Based on the framework proposed by Bottini and Doeller (2020) regarding the use of internal representational spaces to represent non-spatial dimensions, we argue that order coding in the intraparietal sulcus might reflect the use of a low-dimensional space (i.e., a horizontal axis from left to right, as captured in the SPoARC), whereas order coding in the hippocampus might reflect the use of a higher-dimensional space (e.g., an allocentric reference frame).

## Open Practice Statement

This experiment was not preregistered. The data and analysis code supporting the reported findings are available at https://osf.io/3ek6p.

### CRediT authorship contribution statement

The authors’ contributions are reported following the CRediT taxonomy (see https://credit.niso.org/contributor-roles-defined/). **Maëliss Vivion:** Conceptualization, Investigation, Formal analysis, Methodology, Writing – original draft, Writing – review & editing. **Fabien Mathy:** Conceptualization, Funding Acquisition, Methodology, Project administration, Supervision, Writing – review & editing. **Alessandro Guida:** Conceptualization, Funding Acquisition, Methodology, Supervision, Writing – review & editing. **Lydiane Mondot:** Investigation, Project administration. **Stephen Ramanoël:** Conceptualization, Investigation, Funding Acquisition, Methodology, Project administration, Supervision, Writing – review & editing

## Acknowledgments

The authors are deeply grateful for the support of the Neuromod Institute.

## Footnotes

1 Note that although a study by Tian and Fischer-Baum (2025) used both a visual and auditory SPoARC task, the authors did not directly compare the two modalities.

2 The study by Tian and Fischer-Baum (2025) is not included here as it used a resting-state fMRI procedure.

3 On the contrary, the boundary-position models correspond to a treatment of the positions based on saliency (i.e., items at the beginning and end of a sequence typically have a higher perceptual saliency and enhanced distinctiveness compared to middle items, which contributes to their superior recall, Cowan, 2001; Henson, 1998; Page and Norris, 1998).

4 Note that although Tamber-Rosenau et al. (2011), using fMRI, also found a shared network for orienting both external and internal attention (comprising in particular the intraparietal sulcus and superior frontal gyrus), they additionally revealed that these two mechanisms rely on distinct neural activation patterns for external and internal shifts of attention.

5 This interpretation is also consistent with recent findings showing a tendency for mid-sagittal ordinal and number mapping (i.e., a mid-sagittal SNARC effect, Attout et al., 2025; Bourgaux et al., 2023), which could suggest a potential three-dimensional coding of ordinal information.

## References

Abrahamse, E., Van Dijck, J.-P., Majerus, S., & Fias, W. (2014). Finding the answer in space: The mental whiteboard hypothesis on serial order in working memory. Frontiers in Human Neuroscience, 8(932), 1–12. 10.3389/fnhum.2014.00932

Albers, A. M., Kok, P., Toni, I., Dijkerman, H. C., & De Lange, F. P. (2013). Shared representations for working memory and mental imagery in early visual cortex. Current Biology, 23(15), 1427–1431. 10.1016/j.cub.2013.05.065

Amiez, C., & Petrides, M. (2007). Selective involvement of the mid-dorsolateral prefrontal cortex in the coding of the serial order of visual stimuli in working memory. Proceedings of the National Academy of Sciences, 104(34), 13786–13791. 10.1073/pnas.0706220104

Arnott, S., Grady, C., Hevenor, S., Graham, S., & Alain, C. (2005). The functional organization of auditory working memory as revealed by fMRI. Journal of cognitive neuroscience, 17(5), 819–831. 10.1162/0898929053747612

Aronov, D., Nevers, R., & Tank, D. W. (2017). Mapping of a non-spatial dimension by the hippocampal–entorhinal circuit. Nature, 543(7647), 719–722. 10.1038/nature21692

Attout, L., Fias, W., Salmon, E., & Majerus, S. (2014). Common neural substrates for ordinal representation in short-term memory, numerical and alphabetical cognition. PloS one, 9(3), 1–15. 10.1371/journal.pone.0092049

Attout, L., Lefevre, S., & Charras, P. (2025). Mid-sagittal spatialization of working memory: A spoarc effect beyond cultural directionality [SSRN Preprint]. 10.2139/ssrn.5399336

Attout, L., Leroy, N., & Majerus, S. (2022). The neural representation of ordinal information: Domain-specific or domain-general? Cerebral Cortex, 32(6), 1170–1183. 10.1093/cercor/bhab279

Attout, L., Ordonez Magro, L., Szmalec, A., & Majerus, S. (2019). The developmental neural substrates of item and serial order components of verbal working memory. Human brain mapping, 40(5), 1541–1553. 10.1002/hbm.24466

Baddeley, A. (2003). Working memory: Looking back and looking forward. Nature reviews neuroscience, 4(10), 829–839. 10.1038/nrn1201

Bellmund, J. L., Gärdenfors, P., Moser, E. I., & Doeller, C. F. (2018). Navigating cognition: Spatial codes for human thinking. Science, 362(6415), 1–11. 10.1126/science.aat6766

Bicanski, A., & Burgess, N. (2018). A neural-level model of spatial memory and imagery. elife, 7, 1–45. 10.7554/eLife.33752

Bonin, P., Poulin-Charronnat, B., Lukowski Duplessy, H., Bard, P., Vinter, A., Ferrand, L., & Méot, A. (2020). Imabase: A new set of 313 colourised line drawings standardised in french for name agreement, image agreement, conceptual familiarity, age-of-acquisition, and imageability. Quarterly Journal of Experimental Psychology, 73(11), 1862–1878. 10.1177/1747021820932822

Boroditsky, L. (2000). Metaphoric structuring: Understanding time through spatial metaphors. Cognition, 75(1), 1–28. 10.1016/S0010-0277(99)00073-6

Bottini, R., & Doeller, C. (2020). Knowledge across reference frames: Cognitive maps and image spaces. Trends in Cognitive Sciences, 24(8), 606–619. 10.1016/j.tics.2020.05.008

Bottini, R., Mattioni, S., & Collignon, O. (2016). Early blindness alters the spatial organization of verbal working memory. Cortex, 83, 271–279. 10.1016/j.cortex.2016.08.007

Bourgaux, L., de Hevia, M.-D., & Charras, P. (2023). Spatio–numerical mapping in 3D. Experimental Psychology, 70(1), 51–60. 10.1027/1618-3169/a000575

Brett, M., Anton, J.-L., Valabregue, R., & Poline, J.-B. (2002). Region of interest analysis using an spm toolbox [Symposium]. 10.1016/S1053-8119(02)90013-3

Brown, G., Preece, T., & Hulme, C. (2000). Oscillator-based memory for serial order. Psychological review, 107(1), 127–181. 10.1037/0033-295X.107.1.127

Buzsáki, G., McKenzie, S., & Davachi, L. (2022). Neurophysiology of remembering. Annual review of psychology, 73(1), 187–215. 10.1146/annurev-psych-021721-110002

Buzsáki, G., & Tingley, D. (2018). Space and time: The hippocampus as a sequence generator. Trends in cognitive sciences, 22(10), 853–869. 10.1016/j.tics.2018.07.006

Chai, W. J., Abd Hamid, A. I., & Abdullah, J. M. (2018). Working memory from the psychological and neurosciences perspectives: A review. Frontiers in psychology, 9, 1–16. 10.3389/fpsyg.2018.00401

Cona, G., & Semenza, C. (2017). Supplementary motor area as key structure for domain-general sequence processing: A unified account. Neuroscience & Biobehavioral Reviews, 72, 28–42. 10.1016/j.neubiorev.2016.10.033

Cowan, N. (2001). The magical number 4 in short-term memory: A reconsideration of mental storage capacity. Behavioral and Brain Sciences, 24(1), 87–185. 10.1017/S0140525X01003922

Cowan, N., Li, D., Moffitt, A., Becker, T. M., Martin, E. A., Saults, J. S., & Christ, S. E. (2011). A neural region of abstract working memory. Journal of cognitive neuroscience, 23(10), 2852–2863. 10.1162/jocn.2011.21625

Cristoforetti, G., Majerus, S., Sahan, M., Van Dijck, J.-P., & Fias, W. (2022). Neural patterns in parietal cortex and hippocampus distinguish retrieval of start versus end positions in working memory. Journal of Cognitive Neuroscience, 34(7), 1230–1245. 10.1162/jocn_a_01860

Crottaz-Herbette, S., Anagnoson, R., & Menon, V. (2004). Modality effects in verbal working memory: Differential prefrontal and parietal responses to auditory and visual stimuli. Neuroimage, 21(1), 340–351. 10.1016/j.neuroimage.2003.09.019

Davis, T., LaRocque, K. F., Mumford, J. A., Norman, K. A., Wagner, A. D., & Poldrack, R. (2014). What do differences between multi-voxel and univariate analysis mean? how subject-, voxel-, and trial-level variance impact fmri analysis. Neuroimage, 97, 271–283. 10.1016/j.neuroimage.2014.04.037

de Belder, M., Abrahamse, E., Kerckhof, M., Fias, W., & van Dijck, J.-P. (2015). Serial position markers in space: Visuospatial priming of serial order working memory retrieval. PloS one, 10(1), 1–10. 10.1371/journal.pone.0116469

Dehaene, S., Bossini, S., & Giraux, P. (1993). The mental representation of parity and number magnitude. Journal of experimental psychology: General, 122(3), 371–396. 10.1037/0096-3445.122.3.371

D’Esposito, M. (2007). From cognitive to neural models of working memory. Philosophical Transactions of the Royal Society B: Biological Sciences, 362(1481), 761–772. 10.1098/rstb.2007.2086

D’Esposito, M., & Postle, B. R. (2015). The cognitive neuroscience of working memory. Annual review of psychology, 66(1), 115–142. 10.1146/annurev-psych-010814-015031

Dimsdale-Zucker, H., & Ranganath, C. (2018). Representational similarity analyses: A practical guide for functional mri applications. In D. Manahan-Vaughan (Ed.), Handbook of behavioral neuroscience (pp. 509–525, Vol. 28). Elsevier. 10.1016/B978-0-12-812028-6.00027-6

Dolfen, N., Reverberi, S., de Beeck, H. O., King, B. R., & Albouy, G. (2024). The hippocampus represents information about movements in their temporal position in a learned motor sequence. Journal of Neuroscience, 44(37), 1–16. 10.1523/JNEUROSCI.0584-24.2024

Durnez, J., Degryse, J., Moerkerke, B., Seurinck, R., Sochat, V., Poldrack, R. A., & Nichols, T. E. (2016). Power and sample size calculations for fmri studies based on the prevalence of active peaks [BioRxiv Preprint]. https://www.biorxiv.org/content/10.1101/049429v1.full.pdf+html

Fenwick, H., Campitelli, G., & Guida, A. (2025). Spatial organisation in the human mind as a function of the distance between stimuli. Quarterly Journal of Experimental Psychology, 78(6), 1107–1123. 10.1177/17470218241255690

Fias, W., Lammertyn, J., Caessens, B., & Orban, G. (2007). Processing of abstract ordinal knowledge in the horizontal segment of the intraparietal sulcus. Journal of Neuroscience, 27(33), 8952–8956. 10.1523/JNEUROSCI.2076-07.2007

Fias, W., Lammertyn, J., Reynvoet, B., Dupont, P., & Orban, G. (2003). Parietal representation of symbolic and nonsymbolic magnitude. Journal of cognitive neuroscience, 15(1), 47–56. 10.1162/089892903321107819

Franklin, M., & Jonides, J. (2009). Order and magnitude share a common representation in parietal cortex. Journal of cognitive neuroscience, 21(11), 2114–2120. 10.1162/jocn.2008.21181

Fulbright, R., Manson, S., Skudlarski, P., Lacadie, C., & Gore, J. (2003). Quantity determination and the distance effect with letters, numbers, and shapes: A functional MR imaging study of number processing. American journal of neuroradiology, 24(2), 193–200. http://www.ajnr.org/content/24/2/193

Galton, F. (1880). Visualised numerals. Nature, 21(533), 252–256.

Gauthier, B., Prabhu, P., Kotegar, K. A., & van Wassenhove, V. (2020). Hippocampal contribution to ordinal psychological time in the human brain. Journal of Cognitive Neuroscience, 32(11), 2071–2086. 10.1162/jocn_a_01586

Geuter, S., Qi, G., Welsh, R. C., Wager, T. D., & Lindquist, M. A. (2018). Effect size and power in fmri group analysis [Biorxiv Preprint]. https://www.biorxiv.org/content/10.1101/295048v1.full.pdf+html

Gijssels, T., Pitt, B., Bottini, R., Battal, C., Collignon, O., & Casasanto, D. (2019). Modality matters: Space-time mappings differ in vision and audition. Proceedings of the 7th international conference on spatial cognition (rome, italy) [In: Cognitive Processing, Vol. 19, no.1, p. S39]. 10.1007/s10339-018-0884-3

Gillebert, C. R., Mantini, D., Thijs, V., Sunaert, S., Dupont, P., & Vandenberghe, R. (2011). Lesion evidence for the critical role of the intraparietal sulcus in spatial attention. Brain, 134(6), 1694–1709. 10.1093/brain/awr085

Ginsburg, V., Archambeau, K., van Dijck, J.-P., Chetail, F., & Gevers, W. (2017). Coding of serial order in verbal, visual and spatial working memory. Journal of Experimental Psychology: General, 146(5), 632–650. 10.1037/xge0000278

Ginsburg, V., van Dijck, J.-P., Previtali, P., Fias, W., & Gevers, W. (2014). The impact of verbal working memory on number–space associations. *Journal of Experimental Psychology: Learning*, Memory, and Cognition, 40(4), 976–986. 10.1037/a0036378

Guida, A., Abrahamse, E., & van Dijck, J.-P. (2020). About the interplay between internal and external spatial codes in the mind: Implications for serial order. Annals of the New York Academy of Sciences, 1477(1), 20–33. 10.1111/nyas.14341

Guida, A., & Campitelli, G. (2019). Explaining the spoarc and snarc effects with knowledge structures: An expertise account. Psychonomic bulletin & review, 26(2), 434–451. 10.3758/s13423-019-01582-0

Guida, A., Carnet, S., Normandon, M., & Lavielle-Guida, M. (2018). Can spatialisation be extended to episodic memory and open sets? Memory, 26(7), 922–935. 10.1080/09658211.2018.1428350

Guida, A., & Lavielle-Guida, M. (2014). 2011 space odyssey: Spatialization as a mechanism to code order allows a close encounter between memory expertise and classic immediate memory studies. Frontiers in psychology, 5(573), 1–5. 10.3389/fpsyg.2014.00573

Guida, A., Leroux, A., Lavielle-Guida, M., & Noël, Y. (2016). A SPoARC in the dark: Spatialization in verbal immediate memory. Cognitive Science, 40(8), 2108–2121. 10.1111/cogs.12316

Guida, A., Megreya, A. M., Lavielle-Guida, M., Noël, Y., Mathy, F., van Dijck, J.-P., & Abrahamse, E. (2018). Spatialization in working memory is related to literacy and reading direction: Culture “literarily” directs our thoughts. Cognition, 175, 96–100. 10.1016/j.cognition.2018.02.013

Guida, A., & Porret, A. (2022). A SPoARC of music: Musicians spatialize melodies but not all-comers. Cognitive Science, 46(5), 1–15. 10.1111/cogs.13139

Guidali, G., Pisoni, A., Bolognini, N., & Papagno, C. (2019). Keeping order in the brain: The supramarginal gyrus and serial order in short-term memory. Cortex, 119, 89–99. 10.1016/j.cortex.2019.04.009

Haxby, J., Gobbini, I., Furey, M., Ishai, A., Schouten, J., & Pietrini, P. (2001). Distributed and overlapping representations of faces and objects in ventral temporal cortex. Science, 293(5539), 2425–2430. 10.1126/science.1063736

Henson, R. (1998). Short-term memory for serial order: The Start-End model. Cognitive Psychology, 36(2), 73–137. 10.1006/cogp.1998.0685

Henson, R., Burgess, N., & Frith, C. (2000). Recoding, storage, rehearsal and grouping in verbal short-term memory: An fMRI study. Neuropsychologia, 38(4), 426–440. 10.1016/S0028-3932(99)00098-6

Hsieh, L.-T., Gruber, M. J., Jenkins, L. J., & Ranganath, C. (2014). Hippocampal activity patterns carry information about objects in temporal context. Neuron, 81(5), 1165–1178. 10.1016/j.neuron.2014.01.015

Huber, S., Klein, E., Moeller, K., & Willmes, K. (2016). Spatial–numerical and ordinal positional associations coexist in parallel. Frontiers in psychology, 7(438), 1–13. 10.3389/fpsyg.2016.00438

Hurlstone, M. (2019). Functional similarities and differences between the coding of positional information in verbal and spatial short-term order memory. Memory, 27(2), 147–162. 10.1080/09658211.2018.1495235

Ischebeck, A., Heim, S., Siedentopf, C., Zamarian, L., Schocke, M., Kremser, C., Egger, K., Strenge, H., Scheperjans, F., & Delazer, M. (2008). Are numbers special? Comparing the generation of verbal materials from ordered categories (months) to numbers and other categories (animals) in an fMRI study. Human brain mapping, 29(8), 894–909. 10.1002/hbm.20433

Ito, Y., & Hatta, T. (2004). Spatial structure of quantitative representation of numbers: Evidence from the snarc effect. Memory & Cognition, 32, 662–673. 10.3758/BF03195857

Kalm, K., & Norris, D. (2014). The representation of order information in auditory-verbal short-term memory. Journal of Neuroscience, 34(20), 6879–6886. 10.1523/JNEUROSCI.4104-13.2014

Kaminski, M., Brzezicka, A., Kaminski, J., & Blinowska, K. J. (2019). Coupling between brain structures during visual and auditory working memory tasks. International journal of neural systems, 29(3), 1–15. 10.1142/S0129065718500466

Kane, M. J., Hambrick, D. Z., Tuholski, S. W., Wilhelm, O., Payne, T. W., & Engle, R. W. (2004). The generality of working memory capacity: A latent-variable approach to verbal and visuospatial memory span and reasoning. Journal of Experimental Psychology: General, 133(2), 189. 10.1037/0096-3445.133.2.189

Kriegeskorte, N., Mur, M., & Bandettini, P. (2008). Representational similarity analysis-connecting the branches of systems neuroscience. Frontiers in systems neuroscience, 2(4), 1–28. 10.3389/neuro.06.004.2008

Kumar, S., Joseph, S., Gander, P., Barascud, N., Halpern, A., & Griffiths, T. (2016). A brain system for auditory working memory. Journal of Neuroscience, 36(16), 4492–4505. 10.1523/JNEUROSCI.4341-14.2016

Laskay-Horváth, C., Aranyi, G., Pachner, O., Remete, E. P., & Kemény, F. (2025). How is reading related to working memory?—a cross-sectional developmental perspective. Reading Research Quarterly, 60(3), 1–15. 10.1002/rrq.70019

Long, N., & Kahana, M. (2019). Hippocampal contributions to serial-order memory. Hippocampus, 29(3), 252–259. 10.1002/hipo.23025

Maguire, E. A., Burgess, N., Donnett, J. G., Frackowiak, R. S., Frith, C. D., & O’Keefe, J. (1998). Knowing where and getting there: A human navigation network. Science, 280(5365), 921–924. 10.1126/science.280.5365.921

Majerus, S. (2019). Verbal working memory and the phonological buffer: The question of serial order. Cortex, 112, 122–133. 10.1016/j.cortex.2018.04.016

Majerus, S., Attout, L., D’Argembeau, A., Degueldre, C., Fias, W., Maquet, P., Martinez Perez, T., Stawarczyk, D., Salmon, E., Van der Linden, M., Phillips, C., & Balteau, E. (2012). Attention supports verbal short-term memory via competition between dorsal and ventral attention networks. Cerebral Cortex, 22(5), 1086–1097. 10.1093/cercor/bhr174

Majerus, S., Bastin, C., Poncelet, M., Van der Linden, M., Salmon, E., Collette, F., & Maquet, P. (2007). Short-term memory and the left intraparietal sulcus: Focus of attention? further evidence from a face short-term memory paradigm. Neuroimage, 35(1), 353–367. 10.1016/j.neuroimage.2006.12.008

Majerus, S., & Boukebza, C. (2013). Short-term memory for serial order supports vocabulary development: New evidence from a novel word learning paradigm. Journal of experimental child psychology, 116(4), 811–828. 10.1016/j.jecp.2013.07.014

Majerus, S., Cowan, N., Péters, F., Van Calster, L., Phillips, C., & Schrouff, J. (2016). Cross-modal decoding of neural patterns associated with working memory: Evidence for attention-based accounts of working memory. Cerebral Cortex, 26(1), 166–179. 10.1093/cercor/bhu189

Majerus, S., d’Argembeau, A., Martinez Perez, T., Belayachi, S., Van der Linden, M., Collette, F., Salmon, E., Seurinck, R., Fias, W., & Maquet, P. (2010). The commonality of neural networks for verbal and visual short-term memory. Journal of cognitive neuroscience, 22(11), 2570–2593. 10.1162/jocn.2009.21378

Majerus, S., Péters, F., Bouffier, M., Cowan, N., & Phillips, C. (2018). The dorsal attention network reflects both encoding load and top–down control during working memory. Journal of Cognitive Neuroscience, 30(2), 144–159. 10.1162/jocn_a_01195

Majerus, S., Poncelet, M., Greffe, C., & Van der Linden, M. (2006). Relations between vocabulary development and verbal short-term memory: The relative importance of short-term memory for serial order and item information. Journal of experimental child psychology, 93(2), 95–119. 10.1016/j.jecp.2005.07.005

Majerus, S., Poncelet, M., Van der Linden, M., Albouy, G., Salmon, E., Sterpenich, V., Vandewalle, G., Collette, F., & Maquet, P. (2006). The left intraparietal sulcus and verbal short-term memory: Focus of attention or serial order? Neuroimage, 32(2), 880–891. 10.1016/j.neuroimage.2006.03.048

Marshuetz, C., Reuter-Lorenz, P., Smith, E., Jonides, J., & Noll, D. (2006). Working memory for order and the parietal cortex: An event-related functional magnetic resonance imaging study. Neuroscience, 139(1), 311–316. 10.1016/j.neuroscience.2005.04.071

Marshuetz, C., Smith, E., Jonides, J., DeGutis, J., & Chenevert, T. (2000). Order information in working memory: FMRI evidence for parietal and prefrontal mechanisms. Journal of cognitive neuroscience, 12(Supplement 2), 130–144. 10.1162/08989290051137459

Nili, H., Wingfield, C., Walther, A., Su, L., Marslen-Wilson, W., & Kriegeskorte, N. (2014). A toolbox for representational similarity analysis. PLoS computational biology, 10(4), 1–10. 10.1371/journal.pcbi.1003553

Nobre, A., Coull, J., Maquet, P., Frith, C., Vandenberghe, R., & Mesulam, M. (2004). Orienting attention to locations in perceptual versus mental representations. Journal of cognitive neuroscience, 16(3), 363–373. 10.1162/089892904322926700

Oberauer, K. (2005). Binding and inhibition in working memory: Individual and age differences in short-term recognition. Journal of experimental psychology: General, 134(3), 368–387. 10.1037/0096-3445.134.3.368

Oberauer, K. (2009). Design for a working memory. Psychology of learning and motivation, 51, 45–100. 10.1016/S0079-7421(09)51002-X

Oosterhof, N. N., Connolly, A. C., & Haxby, J. V. (2016). CoSMoMVPA: multi-modal multivariate pattern analysis of neuroimaging data in Matlab / GNU Octave. Frontiers in Neuroinformatics, 10(27), 1–27. 10.3389/fninf.2016.00027

Page, M., & Norris, D. (1998). The primacy model: A new model of immediate serial recall. Psychological review, 105(4), 761–781.

Papagno, C., Comi, A., Riva, M., Bizzi, A., Vernice, M., Casarotti, A., Fava, E., & Bello, L. (2017). Mapping the brain network of the phonological loop. Human brain mapping, 38(6), 3011–3024. 10.1002/hbm.23569

Peng, P., Barnes, M., Wang, C., Wang, W., Li, S., Swanson, H. L., Dardick, W., & Tao, S. (2018). A meta-analysis on the relation between reading and working memory. Psychological bulletin, 144(1), 48–76. 10.1037/bul0000124

Popal, H., Wang, Y., & Olson, I. (2019). A guide to representational similarity analysis for social neuroscience. Social cognitive and affective neuroscience, 14(11), 1243–1253. 10.1093/scan/nsz099

Posit team. (2023). Rstudio: Integrated development environment for r. Posit Software, PBC. Boston, MA. http://www.posit.co/

Price, D. (2024). MNI2FS: High resolution surface rendering of mni registered volumes. [matlab toolbox version 1.4.0.0]. *GitHub*. https://www.github.com/dprice80/mni2fs

Protopapa, F., Hayashi, M., Kulashekhar, S., van der Zwaag, W., Battistella, G., Murray, M., Kanai, R., & Bueti, D. (2019). Chronotopic maps in human supplementary motor area. PLoS biology, 17(3), 1–34. 10.1371/journal.pbio.3000026

Rasoulzadeh, V., Sahan, M. I., van Dijck, J.-P., Abrahamse, E., Marzecova, A., Verguts, T., & Fias, W. (2021). Spatial attention in serial order working memory: An EEG study. Cerebral Cortex, 31(5), 2482–2493. 10.1093/cercor/bhaa368

Riggall, A. C., & Postle, B. R. (2012). The relationship between working memory storage and elevated activity as measured with functional magnetic resonance imaging. Journal of Neuroscience, 32(38), 12990–12998. 10.1523/JNEUROSCI.1892-12.2012

Rizza, A., Pedale, T., Mastroberardino, S., Olivetti Belardinelli, M., Van der Lubbe, R., Spence, C., & Santangelo, V. (2024). Working memory maintenance of visual and auditory spatial information relies on supramodal neural codes in the dorsal frontoparietal cortex. Brain Sciences, 14(2), 1–14. 10.3390/brainsci14020123

Roberts, B. M., Libby, L. A., Inhoff, M. C., & Ranganath, C. (2018). Brain activity related to working memory for temporal order and object information. Behavioural brain research, 354, 55–63. 10.1016/j.bbr.2017.05.068

Rodriguez-Jimenez, R., Avila, C., Garcia-Navarro, C., Bagney, A., de Aragon, A. M., Ventura-Campos, N., Martinez-Gras, I., Forn, C., Ponce, G., Rubio, G., et al. (2009). Differential dorsolateral prefrontal cortex activation during a verbal n-back task according to sensory modality. Behavioural brain research, 205(1), 299–302. 10.1016/j.bbr.2009.08.022

Sokolowski, M., Fias, W., Mousa, A., & Ansari, D. (2017). Common and distinct brain regions in both parietal and frontal cortex support symbolic and nonsymbolic number processing in humans: A functional neuroimaging meta-analysis. Neuroimage, 146, 376–394. 10.1016/j.neuroimage.2016.10.028

Stevens, A. A., Skudlarski, P., Gatenby, J. C., & Gore, J. C. (2000). Event-related fmri of auditory and visual oddball tasks. Magnetic resonance imaging, 18(5), 495–502. 10.1016/S0730-725X(00)00128-4

Strick, P., Dum, R., & Fiez, J. (2009). Cerebellum and nonmotor function. Annual review of neuroscience, 32(1), 413–434. 10.1146/annurev.neuro.31.060407.125606

Tamber-Rosenau, B., Esterman, M., Chiu, Y.-C., & Yantis, S. (2011). Cortical mechanisms of cognitive control for shifting attention in vision and working memory. Journal of cognitive neuroscience, 23(10), 2905–2919. 10.1162/jocn.2011.21608

Tian, Y., & Fischer-Baum, S. (2025). The role of spatial processing in verbal serial order working memory. Cognitive, Affective, & Behavioral Neuroscience, 25, 210–239. 10.3758/s13415-024-01240-6

van Dijck, J.-P., Abrahamse, E., Majerus, S., & Fias, W. (2013). Spatial attention interacts with serial-order retrieval from verbal working memory. Psychological science, 24(9), 1854–1859. 10.1177/0956797613479610

van Dijck, J.-P., & Fias, W. (2011). A working memory account for spatial–numerical associations. Cognition, 119(1), 114–119. 10.1016/j.cognition.2010.12.013

Vandenberghe, R., Gitelman, D. R., Parrish, T. B., & Mesulam, M.-M. (2001). Functional specificity of superior parietal mediation of spatial shifting. Neuroimage, 14(3), 661–673. 10.1006/nimg.2001.0860

Vetter, P., Smith, F. W., & Muckli, L. (2014). Decoding sound and imagery content in early visual cortex. Current Biology, 24(11), 1256–1262. 10.1016/j.cub.2014.04.020

Wandell, B. A., Dumoulin, S. O., & Brewer, A. A. (2007). Visual field maps in human cortex. Neuron, 56(2), 366–383. 10.1016/j.neuron.2007.10.012

Weger, U. W., & Pratt, J. (2008). Time flies like an arrow: Space-time compatibility effects suggest the use of a mental timeline. Psychonomic Bulletin & Review, 15(2), 426–430. 10.3758/PBR.15.2.426

Wood, G., Willmes, K., Nuerk, H.-C., & Fischer, M. (2008). On the cognitive link between space and number: A meta-analysis of the SNARC effect. Psychology science quarterly, 50(4), 489–504. https://api.semanticscholar.org/CorpusID:53508874

Yantis, S., Schwarzbach, J., Serences, J. T., Carlson, R. L., Steinmetz, M. A., Pekar, J. J., & Courtney, S. M. (2002). Transient neural activity in human parietal cortex during spatial attention shifts. Nature neuroscience, 5(10), 995–1002. 10.1038/nn921

Zhang, D. R., Li, Z. H., Chen, X. C., Wang, Z. X., Zhang, X. C., Meng, X. M., He, S., & Hu, X. P. (2003). Functional comparison of primacy, middle and recency retrieval in human auditory short-term memory: An event-related fMRI study. Cognitive Brain Research, 16(1), 91–98. 10.1016/S0926-6410(02)00223-9

Zhou, D., Cai, Q., Luo, J., Yi, Z., Li, Y., Seger, C. A., & Chen, Q. (2021). The neural mechanism of spatial-positional association in working memory: A fMRI study. Brain and Cognition, 152, 1–11. 10.1016/j.bandc.2021.105756

Zorzi, M., Di Bono, M. G., & Fias, W. (2011). Distinct representations of numerical and non-numerical order in the human intraparietal sulcus revealed by multivariate pattern recognition. NeuroImage, 56(2), 674–680. 10.1016/j.neuroimage.2010.06.035

