## Supplemental Tables 1-6 for "Distinct cortical regions support the coding of order across visual and auditory working memory"

#### Author Note

**Table S1**

*MNI coordinates for each regions of interest included in the univariate and multivariate analyses*

| Cortical region | Left Hemisphere | Right Hemisphere |
|---|---|---|
| Anterior Intraparietal Sulcus | -38,-41,42 | 44,-31,44 |
| Posterior Intraparietal Sulcus | -28,-61,44 | 28,-59,42 |
| Inferior Frontal Gyrus | -52,9,18 | 40,13,20 |
| Middle Frontal Gyrus | -42,23,30 | 44,33,28 |
| Superior Frontal Gyrus | -22,7,62 | 22,11,58 |
| Anterior Hippocampus | -24,-25,4 | 30,-21,-6 |
| Posterior Hippocampus | -20,-35,0 | 26,-37,0 |
| Supplementary Motor Area | -4,7,56 | |

**Table S2**

*Results of the ANOVA performed on the log-transformed response times*

| Effect | Df | $F$ value | $p$ |
|---|---|---|---|
| Response hand | 1, 39 | 9.44 | .004 |
| Probe position | 2.56, 99.73 | 13.02 | $< .001$ |
| Condition | 1, 39 | 60.01 | $< .001$ |
| Condition $\times$ Probe position | 2.48, 96.86 | 2.53 | .073 |
| Condition $\times$ Response hand | 1, 39 | 0.24 | .630 |
| Response hand $\times$ Probe position | 2.52, 98.31 | 4.62 | .007 |
| Condition $\times$ Response hand $\times$ Probe position | 2.77, 108.11 | 2.61 | .060 |

**Table S3**

*Results of the multivoxel pattern analysis comparing observed representational dissimilarity matrices in regions of interest with four theoretical models*

| Region of interest | Across-modality • Early-late position model | Across-modality • Boundary-position model | Within-modality • Early-late position model | Within-modality • Boundary-position model |
|---|---|---|---|---|
| | | Encoding phase | | |
| IFG | $t(1279) = 4.54$, $p_{FDR} < .001$, $d = 0.69$ | $t(1279) = -1.16$, $p_{FDR} = .291$, $d = -0.18$ | $t(1279) = 9.83$, $p_{FDR} < .001$, $d = 1.54$ | $t(1279) = 5.25$, $p_{FDR} < .001$, $d = 0.82$ |
| MFG | $t(1279) = 4.98$, $p_{FDR} < .001$, $d = 0.78$ | $t(1279) = 2.48$, $p_{FDR} = .021$, $d = 0.39$ | $t(1279) = 3.73$, $p_{FDR} < .001$, $d = 0.58$ | $t(1279) = 1.79$, $p_{FDR} = .099$, $d = 0.28$ |
| SFG | $t(1279) = 3.86$, $p_{FDR} < .001$, $d = 0.60$ | $t(1279) = 2.33$, $p_{FDR} = .030$, $d = 0.36$ | $t(1279) = 3.91$, $p_{FDR} < .001$, $d = 0.61$ | $t(1279) = 2.32$, $p_{FDR} = .030$, $d = 0.36$ |
| IPSa | $t(1279) = 3.41$, $p_{FDR} = .001$, $d = 0.53$ | $t(1279) = 1.01$, $p_{FDR} = .357$, $d = 0.16$ | $t(1279) = 8.43$, $p_{FDR} < .001$, $d = 1.32$ | $t(1279) = 5.94$, $p_{FDR} < .001$, $d = 0.93$ |
| IPSp | $t(1279) = 2.48$, $p_{FDR} = .021$, $d = 0.39$ | $t(1279) = 0.41$, $p_{FDR} = .707$, $d = 0.06$ | $t(1279) = 8.0$, $p_{FDR} < .001$, $d = 1.12$ | $t(1279) = 5.89$, $p_{FDR} < .001$, $d = 0.92$ |
| aHC | $t(1279) = 0.62$, $p_{FDR} = .572$, $d = 0.09$ | $t(1279) = -1.46$, $p_{FDR} = .186$, $d = -0.23$ | $t(1279) = 7.09$, $p_{FDR} < .001$, $d = 1.11$ | $t(1279) = 5.86$, $p_{FDR} < .001$, $d = 0.92$ |
| pHC | $t(1279) = 0.33$, $p_{FDR} = .740$, $d = 0.05$ | $t(1279) = -0.89$, $p_{FDR} = .415$, $d = -0.14$ | $t(1279) = 6.24$, $p_{FDR} < .001$, $d = 0.98$ | $t(1279) = 5.09$, $p_{FDR} < .001$, $d = 0.80$ |
| SMA | $t(1279) = 12.41$, $p_{FDR} < .001$, $d = 1.94$ | $t(1279) = -1.29$, $p_{FDR} = .242$, $d = -0.20$ | $t(1279) = 8.23$, $p_{FDR} < .001$, $d = 1.29$ | $t(1279) = -1.79$, $p_{FDR} = .099$, $d = -0.28$ |
| | | Recognition phase | | |
| IFG | $t(1273) = -2.27$, $p_{FDR} = .032$, $d = -0.07$ | $t(1273) = -2.34$, $p_{FDR} = .029$, $d = -0.08$ | $t(1273) = 8.77$, $p_{FDR} < .001$, $d = 0.29$ | $t(1273) = 8.70$, $p_{FDR} < .001$, $d = 0.28$ |
| MFG | $t(1273) = -2.20$, $p_{FDR} = .036$, $d = -0.09$ | $t(1273) = -2.55$, $p_{FDR} = .018$, $d = -0.08$ | $t(1273) = 7.32$, $p_{FDR} < .001$, $d = 0.24$ | $t(1273) = 7.23$, $p_{FDR} < .001$, $d = 0.24$ |
| SFG | $t(1273) = -1.70$, $p_{FDR} = .099$, $d = -0.06$ | $t(1273) = -1.57$, $p_{FDR} = .123$, $d = -0.05$ | $t(1273) = 4.87$, $p_{FDR} < .001$, $d = 0.16$ | $t(1273) = 4.85$, $p_{FDR} < .001$, $d = 0.16$ |
| IPSa | $t(1273) = -1.53$, $p_{FDR} = .13$, $d = -0.05$ | $t(1273) = -4.10$, $p_{FDR} < .001$, $d = -0.13$ | $t(1273) = 10.32$, $p_{FDR} < .001$, $d = 0.34$ | $t(1273) = 8.36$, $p_{FDR} < .001$, $d = 0.27$ |
| IPSp | $t(1273) = -2.58$, $p_{FDR} = .017$, $d = -0.08$ | $t(1273) = -3.85$, $p_{FDR} < .001$, $d = -0.13$ | $t(1273) = 9.42$, $p_{FDR} < .001$, $d = 0.30$ | $t(1273) = 8.98$, $p_{FDR} < .001$, $d = 0.29$ |
| aHC | $t(1273) = -1.79$, $p_{FDR} = .084$, $d = -0.06$ | $t(1273) = -2.14$, $p_{FDR} = .040$, $d = -0.07$ | $t(1273) = 7.34$, $p_{FDR} < .001$, $d = 0.24$ | $t(1273) = 5.74$, $p_{FDR} < .001$, $d = 0.19$ |
| pHC | $t(1273) = -1.94$, $p_{FDR} = .062$, $d = -0.06$ | $t(1273) = -2.25$, $p_{FDR} = .033$, $d = -0.06$ | $t(1273) = 6.08$, $p_{FDR} < .001$, $d = 0.19$ | $t(1273) = 5.71$, $p_{FDR} < .001$, $d = 0.20$ |
| SMA | $t(1273) = -2.43$, $p_{FDR} = .023$, $d = -0.08$ | $t(1273) = -1.50$, $p_{FDR} = .133$, $d = -0.05$ | $t(1273) = 5.96$, $p_{FDR} < .001$, $d = 0.19$ | $t(1273) = 6.67$, $p_{FDR} < .001$, $d = 0.22$ |

*Note.* Results of post-hoc tests from linear mixed models including theoretical models, ROIs, and their interaction as fixed effects, with subjects as a random effect. The dependent variable was the $z$-transformed correlation between theoretical RDMs and individual neural RDMs. Two models were fit: one for the encoding phase and one for the recognition phase. Reported $p$-values are FDR-corrected using the Benjamini–Hochberg procedure. Effect sizes are reported as Cohen's $d$, computed as the ratio of the estimated marginal mean to the model's residual standard deviation.

### Table S4

*Results of post-hoc pairwise comparisons of theoretical model fits to the neural data within each region of interest*

| Region of interest | Across-modality • Early-late position model *vs.* Across-modality • Boundary-position model | Within-modality • Early-late position model *vs.* Across-modality • Early-late position model | Within-modality • Boundary-position model *vs.* Across-modality • Early-late position model | Within-modality • Early-late position model *vs.* Across-modality • Boundary-position model | Within-modality • Boundary-position model *vs.* Across-modality • Boundary-position model | Within-modality • Early-late position model *vs.* Within-modality • Boundary-position model |
|---|---|---|---|---|---|---|
| Encoding phase | | | | | | |
| IFG | $t(1240) = 3.98$, $p_{FDR} < .001$ Cohen's $d = 0.88$ | $t(1240) = 3.81$, $p_{FDR} < .001$ Cohen's $d = 0.84$ | $t(1240) = 0.57$, $p_{FDR} = .623$ Cohen's $d = 0.13$ | $t(1240) = 7.78$, $p_{FDR} < .001$ Cohen's $d = 1.72$ | $t(1240) = 4.54$, $p_{FDR} < .001$ Cohen's $d = 1.0$ | $t(1240) = 3.24$, $p_{FDR} = .003$ Cohen's $d = 0.72$ |
| MFG | $t(1240) = 1.77$, $p_{FDR} = .335$ Cohen's $d = 0.39$ | $t(1240) = -0.89$, $p_{FDR} = .507$ Cohen's $d = -0.20$ | $t(1240) = -2.26$, $p_{FDR} = .223$ Cohen's $d = -0.50$ | $t(1240) = 0.88$, $p_{FDR} = .507$ Cohen's $d = 0.19$ | $t(1240) = -0.49$, $p_{FDR} = .667$ Cohen's $d = -0.11$ | $t(1240) = 1.37$, $p_{FDR} = .484$ Cohen's $d = 0.30$ |
| SFG | $t(1240) = 1.08$, $p_{FDR} = .490$ Cohen's $d = 0.24$ | $t(1240) = 0.03$, $p_{FDR} = .996$ Cohen's $d = 0.01$ | $t(1240) = -1.09$, $p_{FDR} = .490$ Cohen's $d = -0.24$ | $t(1240) = 1.12$, $p_{FDR} = .490$ Cohen's $d = 0.25$ | $t(1240) = -0.01$, $p_{FDR} = .996$ Cohen's $d = -0.001$ | $t(1240) = 1.12$, $p_{FDR} = .490$ Cohen's $d = 0.25$ |
| IPSa | $t(1240) = 1.70$, $p_{FDR} = .153$ Cohen's $d = 0.37$ | $t(1240) = 3.55$, $p_{FDR} = .002$ Cohen's $d = 0.79$ | $t(1240) = 1.79$, $p_{FDR} = .153$ Cohen's $d = 0.40$ | $t(1240) = 5.26$, $p_{FDR} < .001$ Cohen's $d = 1.16$ | $t(1240) = 3.49$, $p_{FDR} = .002$ Cohen's $d = 0.77$ | $t(1240) = 1.76$, $p_{FDR} = .153$ Cohen's $d = 0.39$ |
| IPSp | $t(1240) = 1.47$, $p_{FDR} = .223$ Cohen's $d = 0.32$ | $t(1240) = 3.91$, $p_{FDR} < .001$ Cohen's $d = 0.86$ | $t(1240) = 2.41$, $p_{FDR} = .046$ Cohen's $d = 0.53$ | $t(1240) = 5.38$, $p_{FDR} < .001$ Cohen's $d = 1.18$ | $t(1240) = 3.88$, $p_{FDR} < .001$ Cohen's $d = 0.86$ | $t(1240) = 1.50$, $p_{FDR} = .223$ Cohen's $d = 0.33$ |
| aHC | $t(1240) = 1.47$, $p_{FDR} = .255$ Cohen's $d = 0.32$ | $t(1240) = 4.58$, $p_{FDR} < .001$ Cohen's $d = 1.01$ | $t(1240) = 3.72$, $p_{FDR} < .001$ Cohen's $d = 0.82$ | $t(1240) = 6.05$, $p_{FDR} < .001$ Cohen's $d = 1.34$ | $t(1240) = 5.19$, $p_{FDR} < .001$ Cohen's $d = 1.15$ | $t(1240) = 0.87$, $p_{FDR} = .490$ Cohen's $d = 0.19$ |
| pHC | $t(1240) = 0.86$, $p_{FDR} = .490$ Cohen's $d = 0.19$ | $t(1240) = 4.19$, $p_{FDR} < .001$ Cohen's $d = 0.92$ | $t(1240) = 3.37$, $p_{FDR} = .003$ Cohen's $d = 0.74$ | $t(1240) = 505$, $p_{FDR} < .001$ Cohen's $d = 1.12$ | $t(1240) = 4.23$, $p_{FDR} < .001$ Cohen's $d = 0.93$ | $t(1240) = 0.82$, $p_{FDR} = .490$ Cohen's $d = 0.18$ |
| SMA | $t(1240) = 9.71$, $p_{FDR} < .001$ Cohen's $d = 2.14$ | $t(1240) = -2.96$, $p_{FDR} = .007$ Cohen's $d = -0.65$ | $t(1240) = -10.06$, $p_{FDR} < .001$ Cohen's $d = -2.22$ | $t(1240) = 6.75$, $p_{FDR} < .001$ Cohen's $d = 1.49$ | $t(1240) = -0.35$, $p_{FDR} = .757$ Cohen's $d = -0.07$ | $t(1240) = 7.10$, $p_{FDR} < .001$ Cohen's $d = 1.57$ |
| Recognition phase | | | | | | |
| IFG | $t(1240) = 0.05$, $p_{FDR} = .988$ Cohen's $d = 0.01$ | $t(1240) = 7.86$, $p_{FDR} < .001$ Cohen's $d = 1.74$ | $t(1240) = 7.81$, $p_{FDR} < .001$ Cohen's $d = 1.73$ | $t(1240) = 7.91$, $p_{FDR} < .001$ Cohen's $d = 1.75$ | $t(1240) = 7.86$, $p_{FDR} < .001$ Cohen's $d = 1.74$ | $t(1240) = 0.05$, $p_{FDR} = .988$ Cohen's $d = 0.01$ |
| MFG | $t(1240) = 0.25$, $p_{FDR} = .988$ Cohen's $d = 0.05$ | $t(1240) = 6.78$, $p_{FDR} < .001$ Cohen's $d = -1.50$ | $t(1240) = 6.71$, $p_{FDR} < .001$ Cohen's $d = -1.48$ | $t(1240) = 7.02$, $p_{FDR} < .001$ Cohen's $d = -1.55$ | $t(1240) = 6.96$, $p_{FDR} < .001$ Cohen's $d = -1.54$ | $t(1240) = 0.06$, $p_{FDR} = .988$ Cohen's $d = 0.01$ |
| SFG | $t(1240) = -0.09$, $p_{FDR} = .988$ Cohen's $d = 0.02$ | $t(1240) = 4.68$, $p_{FDR} < .001$ Cohen's $d = 1.03$ | $t(1240) = 4.66$, $p_{FDR} < .001$ Cohen's $d = 1.03$ | $t(1240) = 4.59$, $p_{FDR} < .001$ Cohen's $d = 1.01$ | $t(1240) = 4.57$, $p_{FDR} < .001$ Cohen's $d = 1.01$ | $t(1240) = 0.02$, $p_{FDR} = .988$ Cohen's $d = 0.003$ |
| IPSa | $t(1240) = 1.83$, $p_{FDR} = .117$ Cohen's $d = 0.40$ | $t(1240) = 8.44$, $p_{FDR} < .001$ Cohen's $d = 1.86$ | $t(1240) = 7.04$, $p_{FDR} < .001$ Cohen's $d = 1.56$ | $t(1240) = 10.27$, $p_{FDR} < .001$ Cohen's $d = 2.27$ | $t(1240) = 8.87$, $p_{FDR} < .001$ Cohen's $d = 1.96$ | $t(1240) = 1.40$, $p_{FDR} = .229$ Cohen's $d = 0.31$ |
| IPSp | $t(1240) = 0.91$, $p_{FDR} = .585$ Cohen's $d = 0.20$ | $t(1240) = 8.54$, $p_{FDR} < .001$ Cohen's $d = 1.89$ | $t(1240) = 8.22$, $p_{FDR} < .001$ Cohen's $d = 1.82$ | $t(1240) = 9.45$, $p_{FDR} < .001$ Cohen's $d = 2.09$ | $t(1240) = 9.14$, $p_{FDR} < .001$ Cohen's $d = 2.02$ | $t(1240) = 0.31$, $p_{FDR} = .929$ Cohen's $d = 0.07$ |
| aHC | $t(1240) = 0.24$, $p_{FDR} = .943$ Cohen's $d = 0.05$ | $t(1240) = 6.50$, $p_{FDR} < .001$ Cohen's $d = 1.44$ | $t(1240) = 5.37$, $p_{FDR} < .001$ Cohen's $d = 1.19$ | $t(1240) = 6.75$, $p_{FDR} < .001$ Cohen's $d = 1.49$ | $t(1240) = 5.61$, $p_{FDR} < .001$ Cohen's $d = 1.24$ | $t(1240) = 1.14$, $p_{FDR} = 460$ Cohen's $d = 0.25$ |
| pHC | $t(1240) = 0.22$, $p_{FDR} = .943$ Cohen's $d = 0.05$ | $t(1240) = 5.71$, $p_{FDR} < .001$ Cohen's $d = 1.26$ | $t(1240) = 5.45$, $p_{FDR} < .001$ Cohen's $d = 1.20$ | $t(1240) = 5.93$, $p_{FDR} < .001$ Cohen's $d = 1.31$ | $t(1240) = 5.67$, $p_{FDR} < .001$ Cohen's $d = 1.25$ | $t(1240) = 0.26$, $p_{FDR} = .943$ Cohen's $d = 0.06$ |
| SMA | $t(1240) = -0.66$, $p_{FDR} = 772$ Cohen's $d = -0.15$ | $t(1240) = 5.97$, $p_{FDR} < .001$ Cohen's $d = 1.32$ | $t(1240) = 6.48$, $p_{FDR} < .001$ Cohen's $d = 1.43$ | $t(1240) = 5.31$, $p_{FDR} < .001$ Cohen's $d = 1.17$ | $t(1240) = 5.82$, $p_{FDR} < .001$ Cohen's $d = 1.29$ | $t(1240) = -0.51$, $p_{FDR} = 772$ Cohen's $d = -0.11$ |

*Note.* Results from pairwise comparisons of estimated marginal means based on linear mixed models including model, ROI, and their interaction as fixed effects, and participant as a random effect. Two models were fit: one for the encoding phase and one for the recognition phase. Reported $p$-values are FDR-corrected using the Benjamini–Hochberg procedure. Effect sizes are reported as Cohen's $d$, reflecting the standardized difference between model predictions.

**Table S5**

*Results of the multivoxel pattern analysis comparing observed representational dissimilarity matrices in regions of interest with two theoretical models, separately for the visual and auditory conditions*

| Region of interest | Auditory condition: Early-late position model | Auditory condition: Boundary-position model | Visual condition: Early-late position model | Visual condition: Boundary-position model |
|---|---|---|---|---|
| | | Encoding phase | | |
| IFG | $t(621) = 2.15$, $p_{FDR} = .103$, $d = 0.34$ | $t(621) = -1.54$, $p_{FDR} = .249$, $d = -0.25$ | $t(640) = 6.12$, $p_{FDR} < .001$, $d = 0.96$ | $t(640) = -1.36$, $p_{FDR} = .352$, $d = -0.21$ |
| MFG | $t(621) = -0.04$, $p_{FDR} = .968$, $d = -0.01$ | $t(621) = 0.41$, $p_{FDR} = .836$, $d = 0.07$ | $t(640) = 2.87$, $p_{FDR} = .014$, $d = 0.45$ | $t(640) = 0.27$, $p_{FDR} = .943$, $d = 0.04$ |
| SFG | $t(621) = 2.37$, $p_{FDR} = .073$, $d = 0.38$ | $t(621) = -0.18$, $p_{FDR} = .912$, $d = -0.03$ | $t(640) = 2.15$, $p_{FDR} = .072$, $d = 0.34$ | $t(640) = 0.73$, $p_{FDR} = .813$, $d = 0.11$ |
| IPSa | $t(621) = 0.45$, $p_{FDR} = .836$, $d = 0.07$ | $t(621) = -0.25$, $p_{FDR} = .912$, $d = -0.04$ | $t(640) = 3.99$, $p_{FDR} < .001$, $d = 0.62$ | $t(640) = 0.66$, $p_{FDR} = .813$, $d = 0.10$ |
| IPSp | $t(621) = 0.97$, $p_{FDR} = .481$, $d = 0.16$ | $t(621) = 1.17$, $p_{FDR} = .434$, $d = 0.19$ | $t(640) = 2.81$, $p_{FDR} = .014$, $d = 0.44$ | $t(640) = -0.34$, $p_{FDR} = .943$, $d = -0.05$ |
| aHC | $t(621) = 1.09$, $p_{FDR} = .444$, $d = 0.17$ | $t(621) = -2.48$, $p_{FDR} = .073$, $d = -0.40$ | $t(640) = 0.15$, $p_{FDR} = .943$, $d = 0.02$ | $t(640) = 0.23$, $p_{FDR} = .943$, $d = 0.04$ |
| pHC | $t(621) = 2.02$, $p_{FDR} = .116$, $d = 0.32$ | $t(621) = -1.58$, $p_{FDR} = .249$, $d = -0.25$ | $t(640) = -0.22$, $p_{FDR} = .943$, $d = -0.03$ | $t(640) = 0.05$, $p_{FDR} = .958$, $d = 0.01$ |
| SMA | $t(621) = 6.42$, $p_{FDR} < .001$, $d = 1.03$ | $t(621) = -3.06$, $p_{FDR} = .019$, $d = -0.49$ | $t(640) = 7.5$, $p_{FDR} < .001$, $d = 1.17$ | $t(640) = -3.02$, $p_{FDR} = .011$, $d = -0.47$ |
| | | Recognition phase | | |
| IFG | $t(640) = 1.21$, $p_{FDR} = .605$, $d = 0.19$ | $t(640) = -0.75$, $p_{FDR} = .657$, $d = -0.12$ | $t(640) = -0.27$, $p_{FDR} = .940$, $d = -0.04$ | $t(640) = 1.38$, $p_{FDR} = .675$, $d = 0.22$ |
| MFG | $t(640) = 1.12$, $p_{FDR} = .604$, $d = 0.17$ | $t(640) = -0.91$, $p_{FDR} = .657$, $d = -0.14$ | $t(640) = -0.11$, $p_{FDR} = .940$, $d = -0.02$ | $t(640) = 0.20$, $p_{FDR} = .940$, $d = 0.03$ |
| SFG | $t(640) = -0.76$, $p_{FDR} = .657$, $d = -0.12$ | $t(640) = 0.25$, $p_{FDR} = .894$, $d = 0.04$ | $t(640) = 0.73$, $p_{FDR} = .929$, $d = 0.11$ | $t(640) = -1.01$, $p_{FDR} = .834$, $d = -0.16$ |
| IPSa | $t(640) = -0.13$, $p_{FDR} = .894$, $d = -0.02$ | $t(640) = -0.77$, $p_{FDR} = .657$, $d = -0.12$ | $t(640) = 1.79$, $p_{FDR} = .675$, $d = 0.28$ | $t(640) = -1.41$, $p_{FDR} = .675$, $d = -0.22$ |
| IPSp | $t(640) = 1.15$, $p_{FDR} = .604$, $d = 0.18$ | $t(640) = -0.39$, $p_{FDR} = .854$, $d = -0.06$ | $t(640) = 1.1$, $p_{FDR} = .834$, $d = 0.17$ | $t(640) = -1.6$, $p_{FDR} = .675$, $d = -0.25$ |
| aHC | $t(640) = 1.80$, $p_{FDR} = .604$, $d = 0.28$ | $t(640) = -1.26$, $p_{FDR} = .604$, $d = -0.19$ | $t(640) = 0.87$, $p_{FDR} = .882$, $d = 0.14$ | $t(640) = -0.08$, $p_{FDR} = .940$, $d = -0.01$ |
| pHC | $t(640) = 1.44$, $p_{FDR} = .604$, $d = 0.23$ | $t(640) = -1.43$, $p_{FDR} = .604$, $d = -0.22$ | $t(640) = -0.31$, $p_{FDR} = .940$, $d = -0.05$ | $t(640) = -0.08$, $p_{FDR} = .940$, $d = -0.01$ |
| SMA | $t(640) = -0.15$, $p_{FDR} = .894$, $d = -0.02$ | $t(640) = 0.58$, $p_{FDR} = .747$, $d = 0.09$ | $t(640) = -0.23$, $p_{FDR} = .940$, $d = -0.04$ | $t(640) = 0.43$, $p_{FDR} = .940$, $d = 0.07$ |

*Note.* Results of post-hoc tests from linear mixed models including theoretical models, ROIs, and their interaction as fixed effects, with subjects as a random effect. The dependent variable was the $z$-transformed correlation between theoretical RDMs and individual neural RDMs. Four models were fit, one for each phase (encoding, recognition) and condition (visual, auditory) combination. Reported $p$-values are FDR-corrected using the Benjamini–Hochberg procedure. Effect sizes are reported as Cohen's $d$, computed as the ratio of the estimated marginal mean to the model's residual standard deviation.

**Table S6**

*Results of post-hoc pairwise comparisons of theoretical model fits to the neural data (Early-late position model > Boundary-position model) within each region of interest, separately for the visual and auditory conditions*

| Region of interest | Auditory condition | Visual condition |
|---|---|---|
| | Encoding phase | |
| IFG | $t(600) = 2.67, p_{FDR} = .020, d = 0.59$ | $t(600) = 5.30, p_{FDR} < .001, d = 1.17$ |
| MFG | $t(600) = -0.33, p_{FDR} = .849, d = -0.07$ | $t(600) = 1.84, p_{FDR} = .105, d = 0.41$ |
| SFG | $t(600) = 1.85, p_{FDR} = .105, d = 0.41$ | $t(600) = 1.01, p_{FDR} = .417, d = 0.22$ |
| IPSa | $t(600) = 0.51, p_{FDR} = .814, d = 0.11$ | $t(600) = 2.36, p_{FDR} = .049, d = 0.52$ |
| IPSp | $t(600) = -0.14, p_{FDR} = .889, d = -0.03$ | $t(600) = 2.24, p_{FDR} = .051, d = 0.49$ |
| aHC | $t(600) = 2.58, p_{FDR} = .020, d = 0.57$ | $t(600) = -0.62, p_{FDR} = .950, d = -0.01$ |
| pHC | $t(600) = 2.60, p_{FDR} = .020, d = 0.58$ | $t(600) = -0.19, p_{FDR} = .950, d = -0.04$ |
| SMA | $t(600) = 6.86, p_{FDR} < .001, d = 1.51$ | $t(600) = 7.46, p_{FDR} < .001, d = 1.65$ |
| | Recognition phase | |
| IFG | $t(600) = 1.39, p_{FDR} = .331, d = 0.31$ | $t(600) = -1.65, p_{FDR} = .489, d = -0.26$ |
| MFG | $t(600) = 1.43, p_{FDR} = .331, d = 0.32$ | $t(600) = -0.22, p_{FDR} = .875, d = -0.05$ |
| SFG | $t(600) = -0.71, p_{FDR} = .634, d = -0.16$ | $t(600) = 1.23, p_{FDR} = .489, d = 0.27$ |
| IPSa | $t(600) = 0.45, p_{FDR} = .655, d = 0.1$ | $t(600) = 2.26, p_{FDR} = .193, d = 0.50$ |
| IPSp | $t(600) = 1.09, p_{FDR} = .439, d = 0.24$ | $t(600) = 1.91, p_{FDR} = .228, d = 0.42$ |
| aHC | $t(600) = 2.17, p_{FDR} = .171, d = 0.48$ | $t(600) = 0.67, p_{FDR} = .808, d = 0.15$ |
| pHC | $t(600) = 2.03, p_{FDR} = .171, d = 0.45$ | $t(600) = -0.16, p_{FDR} = .875, d = 0.03$ |
| SMA | $t(600) = -0.52, p_{FDR} = .655, d = -0.11$ | $t(600) = -0.47, p_{FDR} = .852, d = -0.1$ |

*Note.* Results from pairwise comparisons of estimated marginal means based on linear mixed models including model, ROI, and their interaction as fixed effects, and participant as a random effect. Four models were fit, one for each phase (encoding, recognition) and condition (visual, auditory) combination. Reported $p$-values are FDR-corrected using the Benjamini–Hochberg procedure. Effect sizes are reported as Cohen's $d$, reflecting the standardized difference between model predictions.
